# Telomere length of both parents contributes to heritable POT1 cancer-predisposition syndrome

**DOI:** 10.64898/2026.02.09.704652

**Authors:** Annika Martin, Robert Lu, Alise Blake, Kim E. Nichols, Santiago E. Sanchez, Steven E. Artandi, Sharon A. Savage, Richa Sharma, Dirk Hockemeyer

## Abstract

Germline mutations in *POT1* are linked to familial cancer predisposition, and somatic *POT1* mutations occur recurrently in tumors. These mutations promote oncogenesis by enabling aberrant telomere elongation. For inherited *POT1* mutations, a critical question is the extent to which elongated telomeres are transmitted to the next generation from the *POT1* carrier parent and whether the inherited excessively long telomeres elevate cancer risk. Using a nanopore sequencing approach that provides haplotype-specific telomere length measurements, we examined telomere inheritance in families harboring *POT1* mutations. We found that individuals preferentially inherit their longest telomeres from the carrier parent, consistent with extensive telomere elongation in the carrier germline, whereas comparatively short telomeres are predominantly inherited from the non-carrier parent. Analysis of carrier and non-carrier siblings further showed that telomeres inherited from both parents are elongated in *POT1* carriers, with the shortest telomeres undergoing preferential elongation. These findings support a potential mechanism of genetic anticipation in which *POT1* mutations progressively reduce the likelihood that short telomeres capable of enforcing telomere-based tumor suppression are inherited from the carrier parent. Together, our results demonstrate that telomeres inherited from both parents jointly shape telomere-based tumor suppressive barriers.

**Summary sentence:** Allele specific nanopore sequencing reveals that *POT1* mutations reshape germline and somatic telomere dynamics, uncovering a novel mechanism of generational anticipation driven by preferential elongation of short inherited telomeres.

## Introduction

Mutations in the gene encoding the shelterin protein POT1 occur in both sporadic and familial cancers, including chronic lymphocytic leukemia (CLL), glioma, melanoma, and other malignancies, with an overall estimated prevalence of ∼3% across cancers^1–12^. These mutations promote cancer by driving telomere elongation, bypassing the tumor-suppressive checkpoint normally triggered by critically short telomeres^13^. Yet, how heterozygous *POT1* mutations trigger telomere elongation and give rise to a distinct cancer spectrum, including glioma, melanoma, and CLL, remains poorly understood. POT1, a core shelterin subunit, binds the telomeric single-stranded overhang to limit telomerase-mediated extension and prevent ataxia telangiectasia and Rad3-related (ATR) kinase-mediated DNA damage signaling^14–20^. Germline *POT1* variants cause autosomal dominant cancer predisposition without loss of heterozygosity^1,2,9^ Engineered mutations in human pluripotent and hematopoietic stem cells show that heterozygous *POT1* lead to progressive elongation in cell types with active telomerase while the chromosome end protective function of telomeres remains intact^13^. Notably, rare autosomal recessive POT1 mutations have also been identified in families with telomere biology disorders characterized by short telomeres, including Coats plus syndrome^21–23^.

Germline transmission of mutations that cause pathogenic telomere length changes creates a unique scenario in which not only the mutation itself is transmitted but potentially also its phenotypic consequence, namely an aberrant telomere length in the germline.

Telomere length inheritance is a well-established mechanism of genetic anticipation in telomere biology disorders that arise from short telomeres. Telomere biology disorders are a spectrum of life-threatening conditions that include bone marrow failure, liver disease, lung disease and other complications caused by mutations in telomere maintenance genes that result in short telomeres. In families with these disorders, aberrantly short telomeres inherited from a parent can lead to earlier onset and increased severity of disease symptoms in their children^24–28^. Consistent with this mechanism, decreased generational telomere length following both maternal and paternal inheritance of short telomeres has been documented^27,28^.

It has been recently proposed that offspring of *POT1* mutation carriers who inherit the same germline variant develop cancer several decades earlier due to transmission of long telomeres^4^. However, the underlying mechanisms for this anticipation remain unclear, and bulk telomere length measurement does not consistently detect progressive telomere elongation across consecutive generations^4,5^. This raises two central questions in *POT1* mutation-positive families: First, do descendants of *POT1* mutation carriers inherit aberrantly elongated telomeres? Second, if long telomeres are inherited, is this inheritance sufficient to drive earlier disease onset? Since a small number of critically short telomeres is sufficient to trigger replicative senescence and enforce a proliferative barrier^29–31^, shorter telomeres contributed by the non-carrier parent are expected to preserve tumor suppression.

How inherited long telomeres might overcome this fundamental safeguard, therefore, remains unresolved, as the dynamics of inherited telomere length remain incompletely understood. Oocytes generally possess longer telomeres than sperm, yet sperm telomere length increases with age^32^. After fertilization, zygotic telomere length more closely resembles sperm telomere length, supporting the model that telomeres are “reset” early in development. The mechanism underlying this resetting remains to be determined, but current evidence indicates that it is incomplete, resulting in a complex inheritance pattern with a strong paternal influence^33–36^. Consequently, telomere length is heritable, and individual parental telomeres can differ by more than 6 kb at the zygotic stage^37^.

In *POT1* mutation positive families, where half of the telomeres of each generation derive from the affected parent, it remains unclear (1) whether *POT1* mutations drive excessive telomere elongation in the germline, (2) whether non-carrier telomeres retain tumor suppressive function, and (3) if inheritance of long telomeres causes disease anticipation. To this end, we use Oxford Nanopore long-read sequencing to map haplotype-specific telomere inheritance in families with *POT1* mutations.

## Results

### Complex cancer spectrum in families with germline *POT1* mutations

To assess the impact of *POT1* mutations on telomere length inheritance, we analyzed three unrelated families with germline *POT1* mutations (see Supplementary Table 1 for all samples analyzed in this study).

Proband 1 harbored the previously reported heterozygous *POT1* c.1851_1852del; p.Asp617GlufsTer9 mutation^11,38,39^, which was paternally inherited and is located in the C-terminus of POT1, resulting in truncation of the C-terminal TPP1 binding domain^40^. The proband presented with a posterior fossa ependymoma at age 3 years. Family members available for study included both parents, who were healthy at the time of sample collection.

Proband 2 harbored the previously reported heterozygous *POT1* c.107T>C; p.Tyr36His (Y36H) mutation^10,13^ , which was paternally inherited and is located in the first Oligonucleotide/Oligosaccharide-Binding (OB) fold domain of POT1, a region critical for single-stranded telomeric DNA binding. The proband presented with Hodgkin lymphoma at age 16 years. Family members available for study included both parents and a maternal cousin. At the time of sample collection, the proband’s mother and maternal cousin were healthy, whereas the father, who carried the *POT1* mutation, had previously been diagnosed with testicular cancer.

Proband 3 carried a maternally inherited heterozygous *POT1* c.265_273delinsAATCTT; p.Tyr89_Lys91delinsAsnLeu mutation, which is located in the first OB fold domain of POT1. The proband was diagnosed with Hodgkin lymphoma at age 12 years and was managed with chemotherapy, radiation therapy, and autologous bone marrow transplant. Unfortunately, he experienced disease relapse at 13 and 17 years of age, followed by osteosarcoma at 22 years of age, for which he was treated with chemotherapy and radiation. At 27 years of age, he experienced osteosarcoma relapse and expired at age 29 years. Available family members included his mother, sister, five nephews (his sister’s children), and the nephews’ father, with mutation status confirmed by sequencing. The mother, sister and two of the five nephews were confirmed to harbor the same *POT1* variant as the proband. The proband’s sister was diagnosed with melanoma and CLL at the age of 32 years. Her melanoma was treated with a wide excision, and her CLL was managed with watchful waiting for 8 years before starting therapy on a clinical trial. The proband’s mother was diagnosed with thyroid cancer at 64 years. The nephews, ranging in age from 9 to 16 years, have all been healthy. All *POT1* carriers are followed annually in a cancer predisposition clinic.

In addition, we studied two other unrelated probands with germline *POT1* mutations for whom family members were not available. Specifically, Proband 4 harbored a *POT1* mutation c.1087C>T; p.Arg363Ter and was diagnosed at 2 years of age with high-risk neuroblastoma to which he succumbed at age 5 years. Proband 5 was found to harbor a *POT1* mutation c.1071dup; p.Gln358SerfsTer13 after being diagnosed with embryonal rhabdomyosarcoma of the prostate at 13 years of age. He experienced recurrence at 14 years before expiring at 16 years. Three unrelated control samples from healthy adults, aged 56, 48, and 38 years, were also evaluated (Figure 1A, Supplementary Table 1). These samples served as sequencing controls to assess telomere clustering rather than as controls for comparing telomere length differences across individuals, as discussed below.

**Figure 1:**
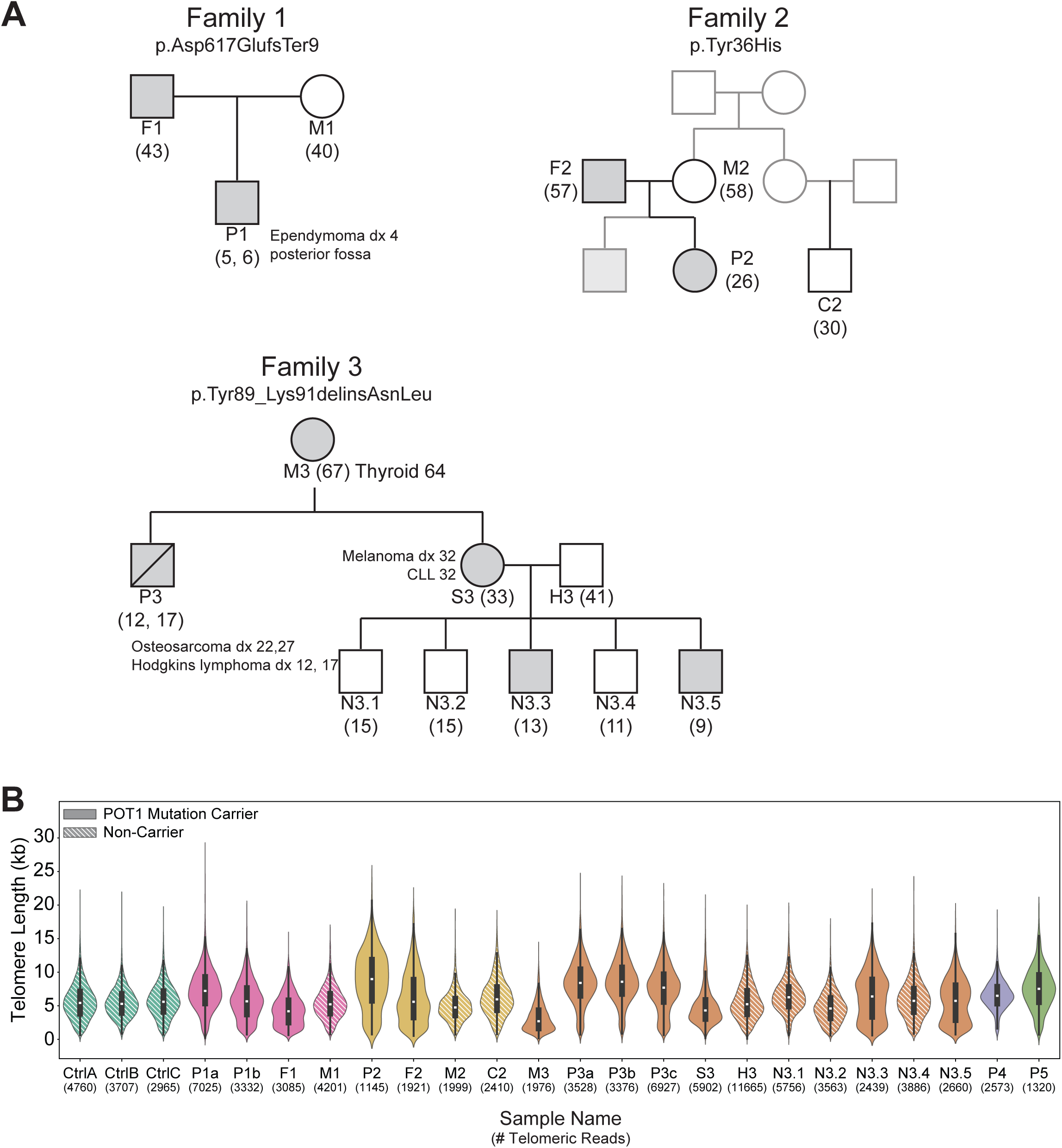
Nanopore long read sequencing of POT1 cancer families. (A) Pedigrees for families with inherited caPOT1 mutations and associated cancer spectrums. Abbreviations as follows: P=Proband, M=Mother, F=Father, S=Sister, H=Husband, N=Nephew. Parenthetical numbers indicate age at time of sample draw (see also Supplementary Table 1). Note for P1 sample was drawn twice at ages 5 and 6 and P3 sample was drawn twice at ages 12 and 17. (B) Bulk telomere length measurements by nanopore sequencing of the indicated individuals. Color groups denote independent families (or unrelated control samples) whereas a hashed fill identifies individuals with no POT1 mutation and solid fill identifies carriers of a caPOT1 mutation.

### Telomere length profiling of *POT1* mutation-positive families by Nanopore sequencing

We applied Nanopore-based telomere-targeted sequencing to evaluate telomere length in this cohort (Supplementary Figure 1)^41–45^. Bulk telomere length measurements closely matched results obtained by telomere restriction fragment (TRF) analysis (Supplementary Figure 2). In total, we sequenced samples from 21 individuals: 11 carrying POT1 mutations (6 individuals with cancer, 3 unaffected carriers), 7 non-carrier family members, and 3 unrelated controls (Figure 1B, Supplementary Table 1). Across samples, 1.3k–12k reads containing telomere ends were detected per individual. Proband samples from Families 1 and 3 were sequenced twice from independent clinical samples, showing strong concordance between replicates, and individual sample measurements were consistent across sequencing rounds (Supplementary Figure 3). Notably, the bulk telomere lengths of 9 POT1 mutation carriers across 3 families, as well as 2 unrelated carriers, ranged widely, from a median length of 4.2 kb to 9.0 kb (Figure 1B). This variability is consistent with previous reports showing that not all POT1 mutation carriers exhibit elongated telomeres and likely reflects differences in allele penetrance, age, disease state, and other contributors to telomere length heterogeneity. These findings highlight the challenges of interpreting average telomere length measurements alone.

### Nanopore sequencing robustly resolves telomere length inheritance patterns in *POT1* families

To trace the inheritance of telomeres across generations, we used a recently published method, Telogator2, to cluster telomere reads based on the Telomere Variant Repeat (TVR) regions located between the telomere and sub-telomere^46^. Extensive polymorphism between TVR regions (Figure 2A) provides haplotype-specific markers for chromosome ends and is subject to Mendelian inheritance. Therefore, while TVRs complicate alignments to the human reference sequence (Supplementary Figure 4), leveraging TVRs to cluster nanopore reads from the same chromosome end in a reference-free manner, as implemented in Telogator2, eliminates the need for a family-specific reference genomes to trace telomere inheritance. By clustering individual reads based on their shared TVR region, we can confidently identify telomeres from single chromosome ends, enabling us to trace their inheritance via comparison to related samples from each family (Figure 2B-D). This method is also robust to minor changes in TVR sequence due to germline expansion/contraction of repeat sequences, a common TVR alteration which contributes to its extensive populational polymorphism (Figure 2D).

**Figure 2:**
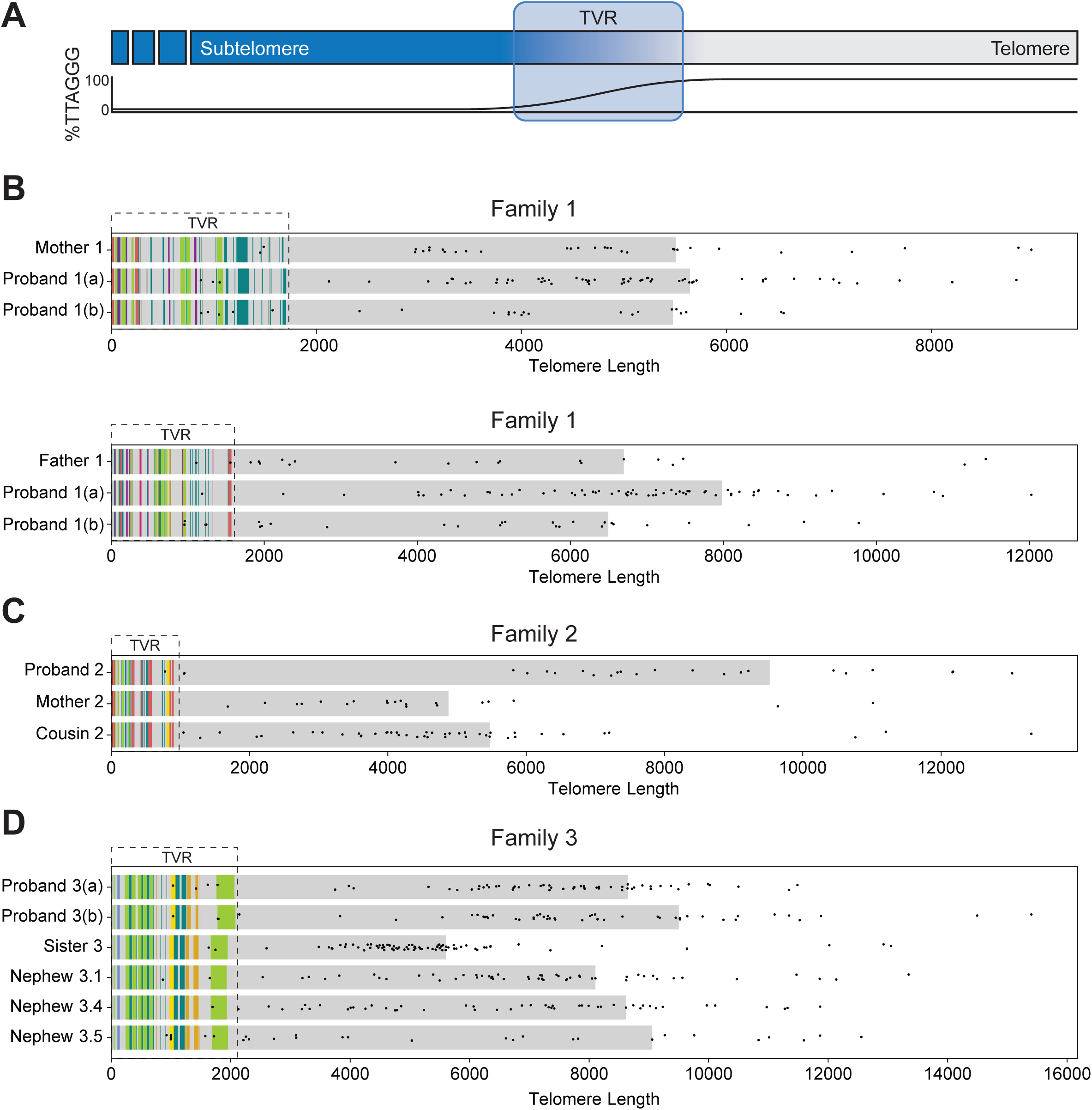
TVR Clustering enables allele correlation between family members. (A) Schematic of the Telomere Variant Repeat (TVR) region situated between canonical TTAGGG repeats of the telomere and the subtelomere comprised of unique chromosomal DNA (B) Example of individual telomere clusters that are shared between Proband 1 and their mother (top) and father (bottom). Individual points show a single measurement of telomere length for the indicated sample, with the grey bar denoting the 75th percentile telomere length for that allele. The TVR region is identified with a dashed line and colored bars within the TVR denote unique telomere variant sequences. For a full table of variant sequences identified and colors used, please refer to the Materials and Methods. (C) As in panel A for an allele identified within patient Family 2 which is shared across the Proband, her Mother, and maternal Cousin (D) As in panel A for an allele identified within patient Family 3 which is shared across six unique DNA samples from patient Family 3.

In total, after filtering out telomere-associated nanopore reads with no detected TVR and potential interstitial alleles, we identified an average of 87 unique telomeric clusters per sample representing 95% of the expected 92 chromosome ends. We then compared clusters from related individuals by aligning TVR consensus sequences using a Levenshtein ratio cutoff of 0.85 (Supplementary Table 2, see Materials and Methods). With this threshold, we found that an average of 97% of TVR haplotypes were identified across independent samples from Proband 1 and Proband 3. In contrast, comparisons between unrelated individuals show only limited haplotype sharing (2%, or 25 of 1068 unique haplotypes across 24 samples) due to the high variability of TVRs among humans. Further, for both Probands 1 and 3, the telomere length of individual haplotype clusters showed strong correlation between two independent sequencings of the same sample and between independent samples from the respective Proband (Supplementary Figure 5). Consistent with original Telogator2 observations^46^, the coefficient of determination (R^2^) for the 75th percentile of telomere length was stronger than either mean or median, likely because it is less affected by outliers e.g. the presence of short telomere fragments generated during sample preparation. Therefore, the 75th percentile telomere length was used for subsequent cluster comparisons^46^.

### The longest telomeres of children with *POT1* mutations are preferentially inherited from their carrier parent

Robust assignment of telomere inheritance allowed us to analyze inheritance patterns in Family 1 and Family 2 with telomere length data from both parents and the proband. Absolute telomere length comparisons are confounded by donor age, cancer presentation, treatment, and sample composition, particularly across generations, due to the significant age differences (see Supplementary Table 1). To overcome these limitations, we rank-ordered telomeres by relative telomere length within each sample, evaluated whether each telomere was inherited from the carrier or non-carrier parent, and compared rank order across family members. This rank-based approach is internally controlled and independent of absolute telomere length, allowing us to isolate the contribution of inherited telomeres and obtain a direct readout of mutation-driven inheritance patterns.

Rank-ordering revealed a significant enrichment of telomeres inherited from the *POT1* mutation-carrier father among the longest telomeres (Figure 3A, B, and Supplementary Figure 6), with a corresponding depletion in the shortest, consistent with germline elongation in the carrier parent. This pattern was not unique to Families 1 and 2; across all assayed individuals with at least one carrier parent sequenced, carrier-derived haplotypes were consistently overrepresented among the longest third of telomeres and underrepresented among the shortest third, highlighting germline elongation as a reproducible, mutation-specific effect (Figure 3C). The shortest telomeres, in contrast, were predominantly inherited from the non-carrier parent, suggesting that these telomeres may still impose functional limits on proliferative capacity even when long telomeres are transmitted from the carrier parent.

**Figure 3:**
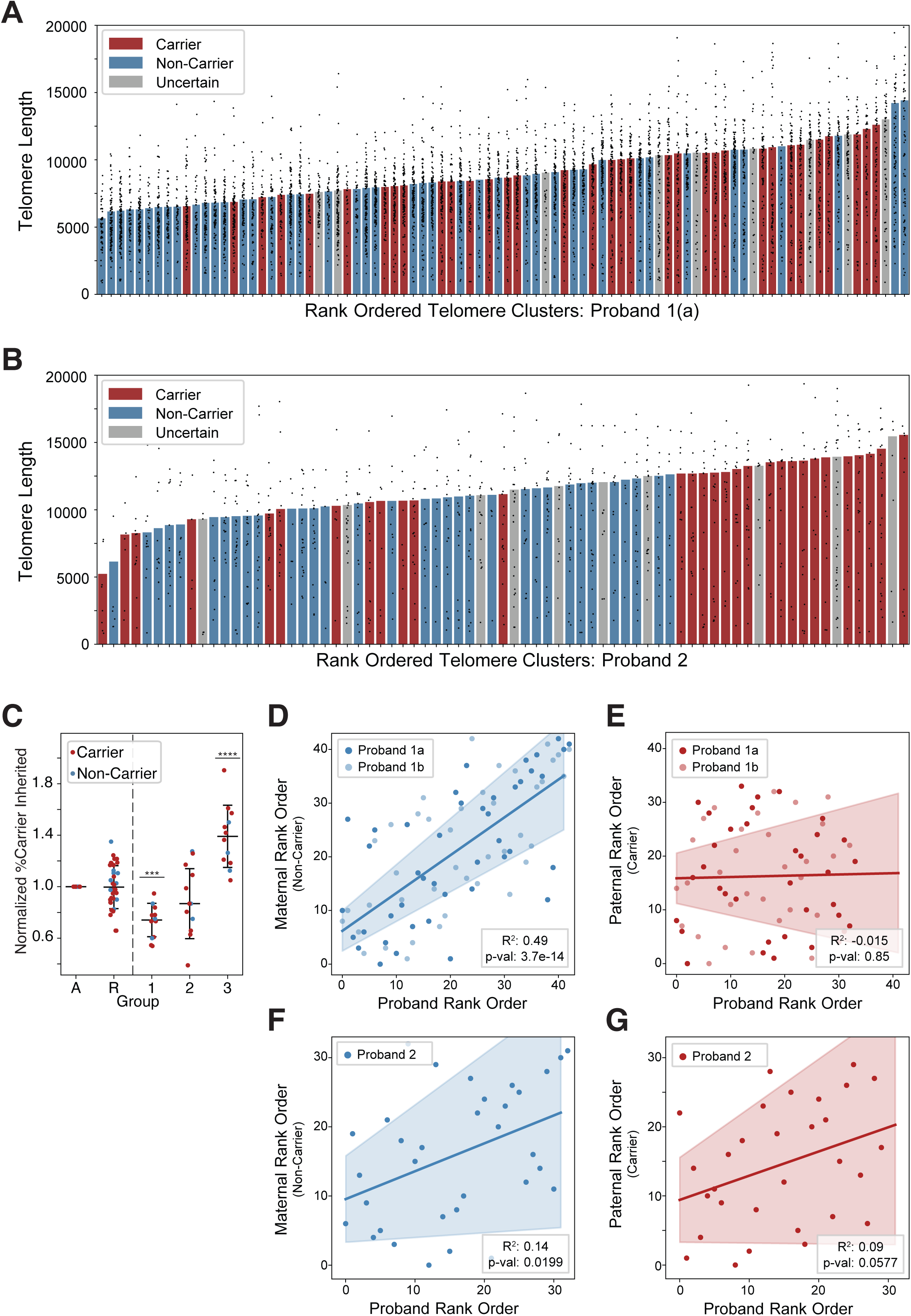
Longest telomeres are preferentially inherited from the caPOT1 mutation carrier parent. (A) All telomere allele clusters identified for Proband 1 rank ordered by 75th percentile telomere length. Points show a single telomere length measurement, whereas bar colors denote whether that allele cluster corresponds to one from the carrier parent (red), the non-carrier parent (blue), or whether the origin could not be identified with our clustering parameters (grey). (B) As in panel A for Proband 2 (C) Percentage of alleles determined to arise from a carrier parent in an indicated fraction normalized to the total percentage of detected carrier alleles: A=All alleles (normalization group), R=random sampling of one third of all telomere alleles for each sample (datapoints represent three independent random sampling events for each sample), 1=shortest third for telomere length for each sample, 2=middle third, and 3=longest third. Each group was compared to the random sampling control using an unpaired t-test with Welch’s correction. Only significant comparisons are shown (*** = p-value<0.001,**** = p-value<0.0001) (D) Shared alleles inherited from the non-carrier parent in Family 1 independently rank-ordered by 75th percentile telomere length for Proband 1 (x-axis) and the non-carrier parent (y-axis). Independent samples for Proband 1 are plotted separately and denoted a and b. Ordinary least squares linear regression was performed using the statsmodels.api OLS module. (E) As in panel D for alleles inherited from the carrier parent. (F) As in panel D for Family 2 (G) As in panel E for Family 2

Direct comparison of Proband 1 rank order with each parent revealed two findings: telomere rank order correlated strongly with the non-carrier mother (Figure 3D) but not with the carrier father (Figure 3E), suggesting that germline elongation “scrambled” the original telomere length order in the carrier. This differential pattern was also observed in absolute haplotype-specific telomere length measurements (Supplementary Figure 7). Together, these results indicate that the excessively long telomeres caused by *POT1* mutations are not fully reset during early embryogenesis. This is consistent with prior studies showing that parental telomere length can be inherited and remain detectable in offspring, reflecting incomplete telomere reprogramming in the zygote^33–36^.

Similar inheritance dynamics were observed in Proband 2, but with notable differences in preservation of parental telomere rank order. Alleles inherited from the non-carrier mother still showed a stronger correlation with the proband’s telomere rank order than those inherited from the carrier father, although this correlation was substantially weaker than in Proband 1 (Figure 4F,G). This reduced preservation of maternal telomere rank order occurred despite a more pronounced enrichment of the longest telomeres among carrier-derived haplotypes in Proband 2 (Figure 4B). These findings suggest that telomere elongation after fertilization progressively disrupts the original inherited telomere-length hierarchy. In this model, telomeres inherited from the carrier parent are already elongated in the germline, whereas the weaker preservation of rank order among telomeres inherited from the non-carrier parent reflects additional post-fertilization elongation acting on both parental haplotypes. The substantially greater loss of parental rank order correlation observed in Family 2 may therefore reflect higher functional penetrance of the POT1 Y36H allele, resulting in more extensive telomere elongation after fertilization and a correspondingly stronger telomere length phenotype.

**Figure 4:**
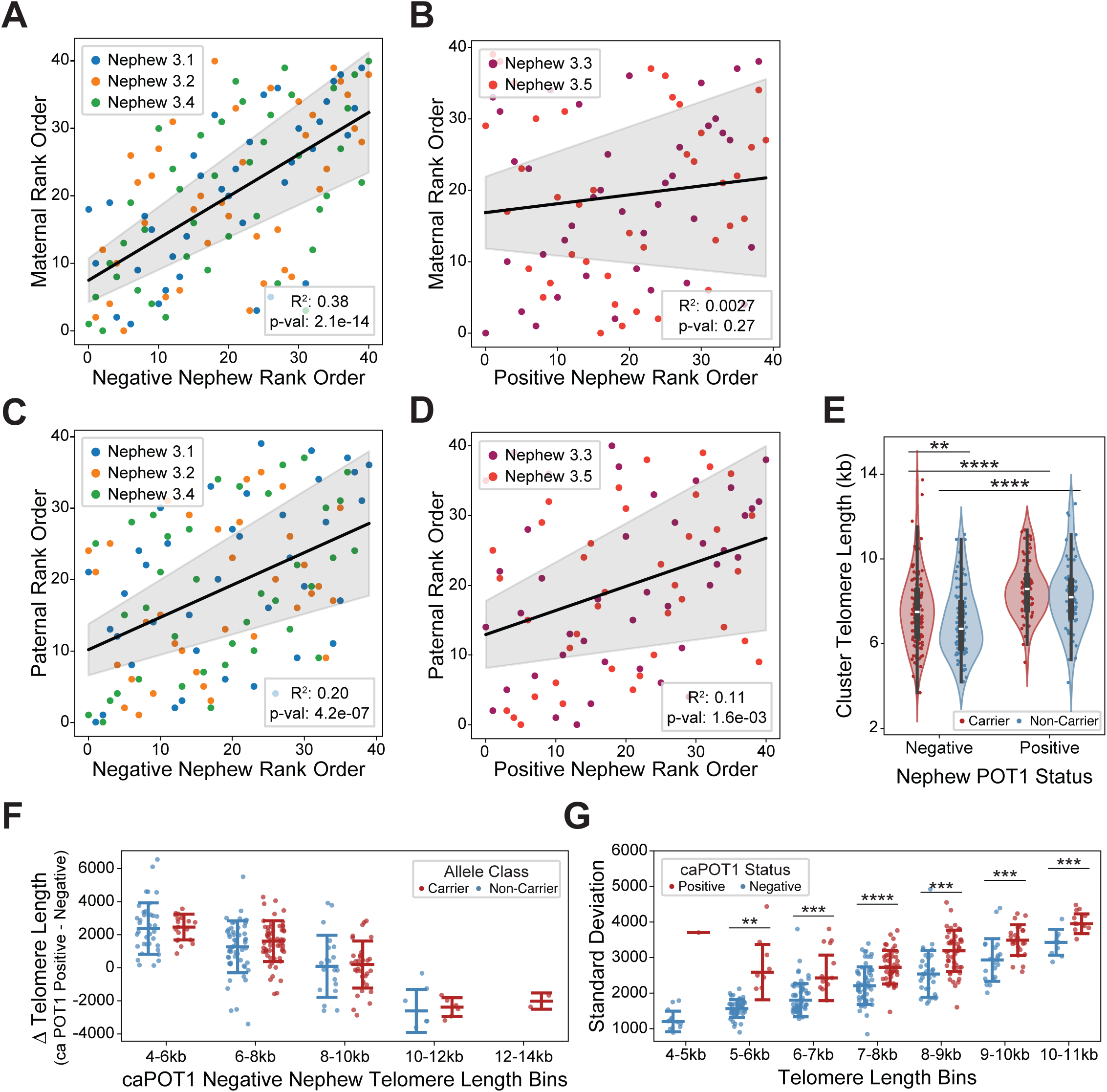
Comparison between carrier and non-carrier children identifies sporadic telomere elongation in POT1 mutation carriers. (A) Shared alleles between Sister 3 and the POT1-negative Nephews independently rank-ordered by 75th percentile telomere length for the Nephews (x-axis) and the Sister (y-axis). Ordinary least squares linear regression was performed using the statsmodels.api OLS module. (B) As in panel A for the caPOT1-positive Nephews. (C) Shared alleles between Husband 3 and the POT1-negative Nephews independently rank-ordered by 75th percentile telomere length for the Nephews (x-axis) and the Husband (y-axis). Ordinary least squares linear regression was performed using the statsmodels.api OLS module. (D) As in panel C for the POT1-positive nephews (E) 75th percentile telomere length for individual telomere allele clusters identified in common between at least two nephews, split into caPOT1-negative (left) and caPOT1-positive (right) nephews and by inheritance from either the carrier parent (red-Sister3) or the non-carrier parent (blue-Husband 3). Group comparisons were performed using Welch’s t-test with FDR-correction for multiple testing. Only significant differences are shown (** = p-value<0.01,**** = p-value<0.0001) (F) Change in 75th percentile telomere length per allele between POT1-positive Nephews and POT1-negative Nephews, binned by allele inheritance (Carrier=red, Non-Carrier=blue) and telomere length in the POT1-negative nephews. No statistical significance by Welch’s t-test between Carrier and Non-Carrier alleles. (G) Standard deviation in telomere length within an individual cluster by binned 75th percentile telomere length for that cluster. All alleles derived from carrier nephews were compared to non-carrier nephews using Welch’s t-test. Only significant p-values are shown (** = p-value<0.01, *** = p-value<0.001,**** = p-value<0.0001)

Finally, the inclusion of the cousin in Family 2, who was of similar age (30 years) to the proband (26 years), enabled us to directly compare the length of telomeres inherited by both individuals from their maternal grandparents. This analysis showed that the shared telomeres in the proband were substantially longer than the corresponding telomeres in the cousin (Supplementary Figure 8) (average 10.5 kb in the proband vs 7.2 kb in the cousin), providing a rough estimate of the absolute telomere elongation that can result from this specific *POT1* mutation.

### A novel mechanism of genetic anticipation through biased telomere elongation

Because inter-allele comparisons are complicated by allele penetrance, we next analyzed Family 3, comparing telomeres from the two nephews who harbor the maternally inherited *POT1* variant with those of their three non-carrier siblings. Rank-order comparison revealed that telomere length in the unaffected nephews correlated strongly with that of their mother (Figure 4A), whereas this correlation was lost in the affected nephews (Figure 4B). Similar to Families 1 and 2, these findings indicate that telomere length patterns inherited from the carrier parent become progressively altered upon transmission to mutation-positive offspring. In contrast, telomeres inherited from the non-carrier father retained a statistically significant rank-order correlation in both carrier and non-carrier nephews (Figure 4C and 4D). However, this correlation showed substantially greater variance in the *POT1*-positive nephews, suggesting that the inherited paternal telomere hierarchy was also partially disrupted in the presence of the mutation. Together, these findings support a two-stage model of telomere remodeling in *POT1* mutation carriers: an initial alteration of telomere length occurring in the carrier parent’s germline, followed by additional post-fertilization telomere elongation in mutation-positive offspring that progressively disrupts the inherited telomere rank order across both parental haplotypes.

To explore post-fertilization dynamics, we directly compared absolute telomere lengths between carrier and non-carrier nephews, who are similar in age, share the same germline origin, and lack cancer or hematopoietic malignancies. In non-carrier nephews, haplotypes from the carrier parent were significantly longer than those from the non-carrier parent (median of haplotype-specific telomere lengths 7.5kb vs 6.7kb), indicating that long telomeres inherited from carriers persist through development without being fully reset (Figure 4E). By contrast, in *POT1* mutation-positive nephews, there was no significant telomere length difference between parental carrier versus non-carrier haplotypes (Figure 4E). This suggests that *POT1* mutations strongly exacerbate post-fertilization telomerase-mediated elongation of the shortest telomeres^26^, inherited from the non-carrier in the next generation. To directly test this hypothesis, we compared telomere length differences between *POT1* mutation-positive nephews and their non-carrier siblings. For each telomere haplotype shared between carrier and non-carrier siblings, we first calculated the 75th percentile telomere length of all reads derived from either a *POT1* mutation-positive or *POT1* mutation-negative nephew. The difference in these observed telomere lengths represents the relative post-fertilization elongation at a per-allele level. When binned by the telomere length of the “baseline” *POT1* mutation-negative nephews, we see robust elongation of the shortest telomeres, irrespective of the parental origin of the individual haplotype (Figure 4F). This indicates that cancer-associated *POT1* mutations generally preserve the tendency of telomerase to act on the shortest telomeres, while increasing overall telomerase activity at telomere ends. However, elongation was not confined to short ends; telomeres across all length classes exhibited sporadic extension, revealing a stochastic component of *POT1* mutation-driven telomere elongation that contributes to increased telomere length heterogeneity in *POT1* mutation carriers (Figure 4G). These findings suggest that generational telomere elongation in *POT1* mutation-positive families is not uniform, but instead reflects a progressive reduction in the probability of inheriting protectively short telomeres from the carrier parent. Consequently, the burden of transmitting short telomeres to maintain telomere-based tumor suppressive barriers increasingly shifts to the non-carrier parent. This model provides a potential explanation for the variable penetrance and heterogeneous anticipation observed in POT1-associated disease.

### Short telomeres remain functionally relevant in a cancer-context-dependent manner

To investigate the role of these inherited alleles in a cancer context, we compared telomeres of the proband and his sister in Family 3, who both carry the *POT1* mutation and share ∼50% of their genetic information but presented with different hematopoietic cancers (HL versus CLL) at 12 and 32 years of age, respectively. Bulk measurements revealed striking differences: the proband had very long telomeres in bone marrow samples that were free of cancer cells at the time of collection (median 8.5 kb), whereas his sister had significantly shorter telomeres (median 4.3 kb) in a peripheral blood sample obtained after her CLL diagnosis (Figure 1B). These differences underscore the challenges of interpreting absolute telomere length across individuals with distinct disease courses, age, treatments, and cellular compositions.

Despite these dramatic differences in absolute telomere length, rank-order analysis revealed a clear hereditary signature, with haplotypes from the *POT1* carrier mother enriched in the longest third of telomeres and depleted in the shortest third in both siblings (Figures 5A and B, Supplementary Figure 6). Rank-order comparison between the siblings’ shared haplotypes also showed significant correlation (Figure 5C). Notably, the sister’s shortest chromosome-arm telomeres displayed constrained heterogeneity both relative to her longest telomeres and compared to her proband brother (Figure 5D).

**Figure 5:**
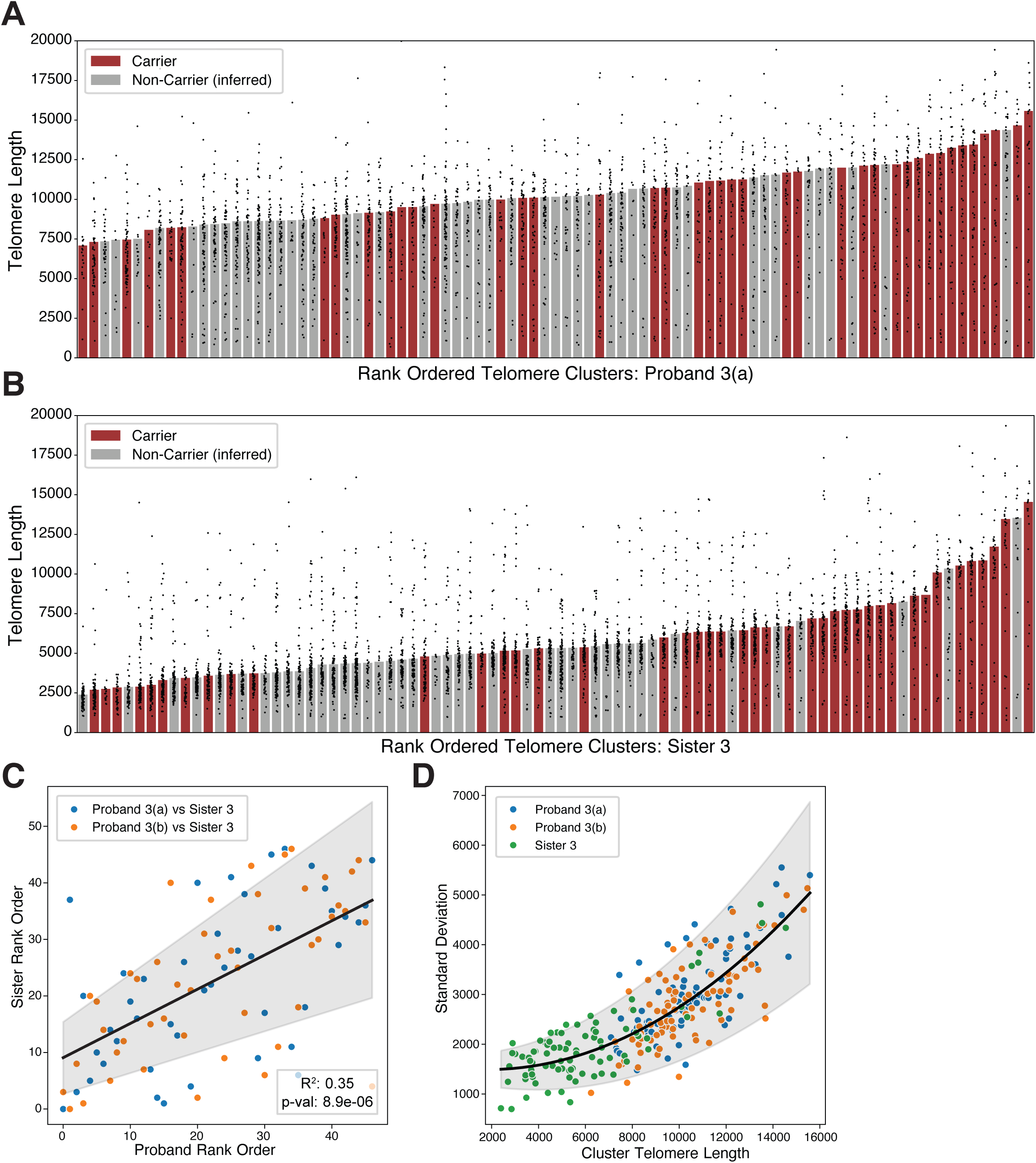
Cancer presentation significantly alters telomere length profile in carrier siblings. (A) All telomere allele clusters identified for Proband 3 rank ordered by 75th percentile telomere length. Points show a single telomere length measurement, whereas bar colors denote whether that allele cluster corresponds to one from the carrier parent (red) or whether the origin could not be confidently identified because there is no sample from the non-carrier parent (grey). (B) As in panel A for Sister 3. (C) Shared alleles between Proband 3 and Sister 3 independently rank-ordered by 75th percentile telomere length for the Proband (x-axis) and the Sister (y-axis). Ordinary least squares linear regression was performed using the statsmodels.api OLS module. (D) Standard deviation in length for individual telomere clusters plotted against 75th percentile telomere length for both Proband 2 samples and Sister 2. Quadratic regression was performed using the scipy.optimize curve_fit module, resulting in the following equation: 1.88e-5 * x2 - 0.068 * x + 1552.55. The grey region indicates the confidence interval defined by the square root of the covariance matrix.

Together, these findings suggest that despite inheriting long telomeres from the carrier parent, short telomeres remain functionally relevant under conditions of high replicative stress, such as CLL. Importantly, 3 of the 5 shortest telomeres in the sister were inherited from the carrier parent. Thus, while telomeres inherited from the carrier parent preferentially contribute to the longest telomeres in the next generation, inheritance of short telomeres from both parents continues to shape the lower end of the telomere length distribution. These observations suggest that telomere-mediated tumor suppressive barriers are influenced not only by POT1-driven telomere elongation but also by the inherited reservoir of short telomeres contributed by both parental haplotypes.

## Discussion

*POT1* mutations promote cancer by elongating telomeres in a telomerase-dependent manner ^12,13,15,47^. Telomerase expression is tightly regulated during development and across cell types, creating contexts in which *POT1* mutations can facilitate tumorigenesis. Early telomerase expression ensures that telomeres are sufficient for proliferation; later restricted activity maintains replicative potential in adult stem cells, and aberrant reactivation in cancers allows bypass of senescence. In germ cells, telomerase preserves telomere length across generations. Understanding how abnormal telomere changes are inherited has been difficult due to limited samples and a lack of haplotype-specific resolution. Nanopore-based chromosome arm-specific measurements now make this possible, and we applied this approach to measure telomere length in families with cancer-associated *POT1* mutations.

Our data show that telomeres inherited from POT1 carrier parents are consistently enriched among the longest third of haplotype-grouped telomeres, while short telomeres are preferentially inherited from the non-carrier parent. This pattern was preserved across both families, indicating that germline elongation in carriers is counteracted by persistence of short non-carrier haplotypes. These findings argue against a simple model of genetic anticipation, as the rank order of short telomeres is preserved and, in some cases, such as the sister of proband 2 with CLL, telomeres eroded to the point of constrained heterogeneity characteristic of CLL^9,48,49^.

While our study was not designed to establish clinical evidence for genetic anticipation and additional supporting evidence is needed, our findings provide a potential mechanistic explanation for how variable anticipation, incomplete penetrance, and heterogeneous disease presentation could arise among carriers of the same pathogenic POT1 allele. Our data suggest a model in which *POT1* mutations do not uniformly elongate telomeres across all chromosome ends or generations. Instead, *POT1* mutations progressively reduce the likelihood that a carrier parent transmits protectively short telomeres to the next generation that are capable of mediating telomere-based tumor suppression (Figure 6), thereby providing a potential mechanism for anticipation. Because telomerase preferentially acts on the shortest telomeres, this increasing proportion of long telomeres may also divert more telomerase activity to the shortest telomeres than would occur if all zygotic telomeres started at comparable lengths. This may, in turn, result in more dramatic telomere elongation of the protective telomere population, despite the re-introduction of short telomeres in each generation through inheritance from the non-carrier parent, Therefore, the cancer risk associated with inheritance of long telomeres appears to be governed by two stochastic processes: the stochastic elongation observed in *POT1* mutation carriers, and the abundance and distribution of the shortest telomeres inherited from both parents within an individual genome. Consistent with this model, our data show that telomeres originating from a carrier parent can still contribute to the pool of the shortest telomeres. The mechanism described above also offers a potential explanation for a long-standing unresolved question: why individuals with elongated telomeres, even in the absence of *POT1* mutations, show increased cancer risk^50–53^. This has been difficult to reconcile with the fact that tumor suppression is enforced by only a small subset of the shortest telomeres^29–31^. We propose that long telomeres are not directly oncogenic. Instead, their presence, even without *POT1* mutations, may reflect depletion of the protectively short telomeres required to enforce telomere-based tumor suppression.

**Figure 6:**
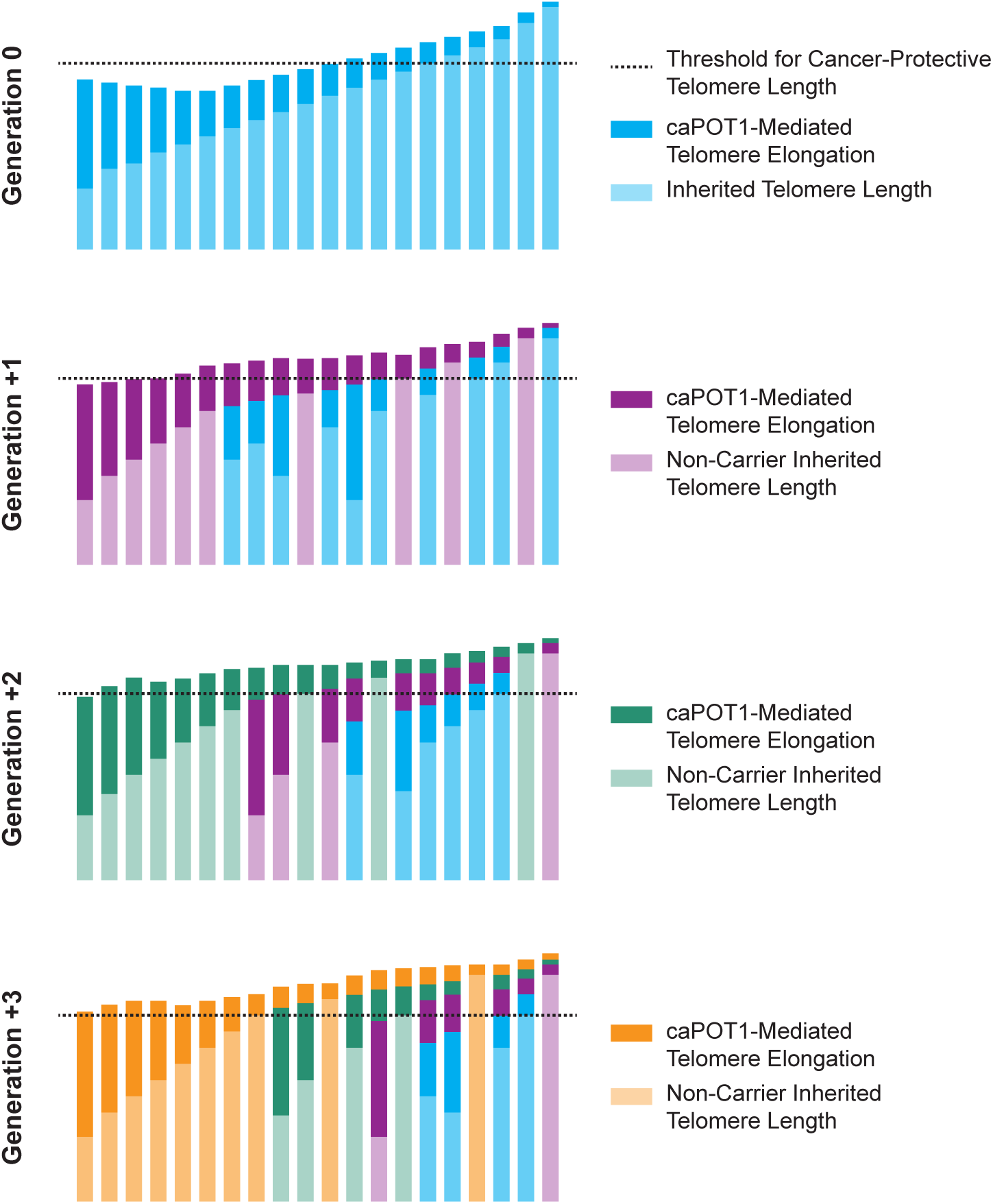
A model for generational anticipation via germline transmission of elongated telomeres. Schematic of a possible mechanism by which POT1 alters disease risk across generations. In the first generation carrying a germline POT1 mutation, telomeres become elongated, with the shortest telomeres being preferentially extended. This over-elongation causes the telomeres in the next generation (G1) that remain sufficiently short to exert tumor-suppressive effects to be more likely inherited from the non-carrier parent. Telomeres are then elongated again in the germ cells of the G1 carrier. Because short telomeres are preferentially extended, the overall number of telomere alleles below the threshold required for tumor suppression decreases further. Consequently, the number of telomeres inherited from the carrier parent that retain the potential to contribute to tumor suppression progressively declines across generations. Over time, this shifts the burden of tumor-suppressive short telomeres almost exclusively to the non-carrier parent. As the proportion of long telomeres increases, this may also result in proportionately more elongation of the shortest telomeres. While this schematic shows only a single representation, it should be noted that the dynamics of telomere elongation and generational inheritance will vary widely by allele penetrance.

This study is constrained by the limited number of families and incomplete sample banking across family members, reflecting the relative rarity of inherited *POT1* mutations. Larger cohorts will be needed to define more precisely how telomere inheritance patterns are transmitted across generations. Our data further suggest that comparing telomere inheritance patterns across families carrying different *POT1* mutations will likely be challenging, as the degree of germline and post-fertilization telomere elongation may depend on the penetrance of individual *POT1* alleles. This is consistent with the diverse and complex tumor spectrum associated with *POT1* mutation that remains a subject of ongoing debate.

Nonetheless, our findings highlight that long telomeres elongated in the germline of *POT1* mutation carriers are transmitted to the next generation, while short telomeres, predominantly from the non-carrier parent, persist and remain functionally relevant. Thus, inheritance of the shortest telomeres from both parents may represent a key determinant of cancer predisposition in POT1-associated disease.

## Acknowledgements

We are grateful to the study participants and their families for their valuable contributions to this study.

We thank the members of the Hockemeyer lab for advice and critical comments on the manuscript, Daniel Rokhsar and Alison Bertuch for critical discussions on this project and comments on the manuscript. We thank Peter Baumann and Nathaniel Deimler for advice and support during the early stage of the project. We also acknowledge QB3 Genomics, UC Berkeley, Berkeley, CA (RRID:SCR_022170) for supplying the sequencing equipment/computational resources for Nanopore sequencing. The Hockemeyer lab is supported by the Innovative Genomics Institute, Siebel Stem Cell Institute the American Cancer Society (133396-RSG-19-029-01-DMC) and the NIH R01HL131744-05 and R21AG095841-01.

This research included experiments conducted by the St. Jude Biorepository which is supported by American Lebanese Syrian Associated Charities. We would like to thank the St. Jude Children’s Research Hospital Biorepository team for their efforts. The work of S.A.S was supported by the Intramural Research Program of the National Institutes of Health (NIH). The contributions of the NIH author(s) are considered Works of the United States Government. The findings and conclusions presented in this paper are those of the author(s) and do not necessarily reflect the views of the NIH or the U.S. Department of Health and Human Services.

## Data availability

Nanopore sequencing fastq reads generated in this study are available at NCBI’s Sequence Read Archive (SRA) and is available through the accession number PRJNA1420205 (https://dataview.ncbi.nlm.nih.gov/object/PRJNA1420205?reviewer=q1fq123cpnri2jv3e1b9o67a <u>pc</u>). Remaining source data is provided with this paper.

## Code availability

The Telogator2 package used for TVR and telomere read analysis of Nanopore reads is available at Github (https://github.com/zstephens/telogator2). Jupyter notebook and python code used for downstream analysis is available on Github (https://github.com/Hockemeyer-Lab/PublishedCode/tree/main/POT1_Telomere_Length_Inheritance). The accompanying PatientKey Excel file used for this analysis is in Supplementary Data 2.

## Author contributions

A.M, R.S, A.B., and D.H. conceptualized the project. A.B. and R.S. and K.E.N, obtained the patient samples. A.M. and R.L. and D.H. designed the experiments. R.L. performed the nanopore sequencing experiments with the support of S.E.S and S.E.A. A.M. analyzed the data. All authors contributed to writing the manuscript.

## Material and Methods

### Patient sample collection and DNA isolation

The study was conducted in accordance with the US Common Rule and was deemed exempt by the Institutional Review Board at St. Jude Children’s Research Hospital (SJCRH; IRB Number: 24-1642, Study Number: POT1). Patients with POT1 tumor predisposition syndrome who underwent genetic counseling and germline genetic testing in the genetic predisposition clinic at SJCRH were included in this study. Demographic, clinical, family history, and germline genetic test results were collected through a review of electronic medical records of patients and families affected by POT1 Tumor Predisposition. Due to the retrospective nature of the study, written informed consent was not required. Note that for Proband 1, two peripheral blood samples were collected at clinic visits separated by four months. For Proband 3, bone marrow samples were collected and banked at age 12, shortly after HL diagnosis and prior to initiating radiation and chemotherapy, and again at age 17 at the time of HL recurrence. Both bone marrow samples were negative for malignant cells. The peripheral blood sample from proband 3’s sister was collected and banked at age 33 following her diagnosis with CLL.

Germline samples were collected and banked in the SJCRH biorepository under the SJFAMILY (NCT03050268) and/or TBANK (NCT01354002) studies. Patient family members were enrolled in SJFAMILY, a study that aims to learn more about the genetic causes of cancer. Patients were enrolled in SJFAMILY and/or TBANK, a study that aims to provide a biorepository of tumor and germline samples for research purposes. Through SJFAMILY and/or TBANK, germline samples were collected and banked for study participants. Samples consisted of blood or healthy bone marrow, aliquoted as cell suspensions or extracted DNA. Primary investigators on both protocols provided permission to utilize the samples in this study. An application was submitted and approved by the SJCRH Tissue Resource Distribution Committee. A Tissue Use Agreement was provided and a Materials Transfer Agreement was executed so the samples could be shared with the Hockemeyer Lab.

Samples for patient Family 2 were obtained through participation in a longitudinal cohort study of individuals and families at high risk of hematologic cancer (ClinicalTrials.gov Identifier NCT00039676). All samples used in this study were de-identified.

### Telomere length nanopore sequencing

Nanopore libraries were prepared following modified library preparation and enrichment protocols previously described by others ^41–45,54^. Specifically, 15-20 ug of purified gDNA were ligated to 2uM of Telobait (Supplementary Table 3) at 35C overnight with 4000 U of T4 DNA ligase in 200-300 uL reaction volume. Ligation reactions were then heat inactivated at 65C for 10 min followed by digestion with 5uL EcoRI-HF . Subsequently, digested DNA samples and ligation ligation reaction were cleaned up with 0.5X ratio of DNA size selection beads (AMPure XP alternative) from the UC Berkeley DNA Sequencing Facility (https://ucberkeleydnasequencing.com/dna-storeroom). Telobait-ligated DNA was resuspended in 100uL Milli-Q water (MQW) and enriched with 60uL of M280 Streptavidin Dynabeads incubation at RT overnight. Next day, beads were washed twice with 1X Binding and Wash (B&W) buffer (prepared from 2X: 10 mM Tris-HCl pH 7.5, 1 mM EDTA, 2 M NaCl as recommended by manufacturer, Thermo Scientific) , equilibrated with 1X Cutsmart buffer for 5 min (without disturbing the beads) then subject to on-bead incubation with 5U T4 DNA polymerase (NEB) fill-in for 30 min at 37C. Beads were then washed again twice with 1X B&W buffer, equilibrated with Cutsmart and resuspended in 100uL of Cutsmart+EcoRV-HF (5uL) to elute the ligated Telobait for at least 6 h at 37C. Eluted Telobait fragments were purified with 1.1X Pronex beads and allowed to dry for at least 10 minutes up to 30 minutes prior to elution ensuring no residual ethanol carry-over. Eluted DNA was quantitated using the Qubit-HS dsDNA kit. Telobait fragments were end-prepped for Nanopore sequencing according to the Native Barcoding Kit 24 V14 (SQK-NBD114-24) protocol up to 14 sample barcodes with the following key modifications. For DNA-cleanup steps, 1.1X relative sample volume of Pronex beads were used instead of AMPure XP beads except for the final cleanup post-NA sequencing adapter ligation whereby 0.5X volume of AMPureXP beads were used instead. Bead drying steps were also extended to 10-15 minutes for cleaning up EcoRV eluates and after end-prep steps. Notably, drying was extended to 30-60 minutes when cleaning-up pooled barcoded Telobait fragments, in order to ensure no residual ethanol carryover prior to NA sequencing adapter ligation. Telobait fragments were eluted in EB buffer according to manufacturer’s protocol and the entire library loaded on the Promethion (R10.4.1) flow cell. Sequencing runs were performed up to 72 h with SUP (superaccuracy) basecalling with a FASTQ quality score cutoff > 8, min read length filter of 500 bp, barcode demultiplexing, with trimming corresponding to the Native Barcoding Kit 24 V14 (SQK-NBD114-24).

### Telomere length restriction fragment analysis

Genomic DNA was prepared as described previously. Briefly, genomic DNA was digested with MboI, AluI, and RNase A overnight at 37°C. The resulting DNA was normalized, and 2 µg of digested DNA was resolved via pulse-field gel electrophoresis and subject to in-gel hybridization with radiolabelled telomeric probe as described previously^20^. Gels were washed three times for 10 min in 0.2× SSC at room temperature then exposed on a phosphorimager screen. Exposed TRF images were analyzed for telomere length as described previously^55^. Analogous to binned lane intensities, phosphorimager gels were analyzed with Fiji/ImageJ2^56^ using the Plot Profile tool for each lane starting from the well to the end of the gel to generate pixel-distance lane intensity profiles. The non-linear regression function of MW to pixel distance was fitted to the below equation model: a+b/(1+(x/c)^d) where x is the distance from the well and y is the DNA MW in (kb) and was performed using MyCurveFit.com (MyAssays Ltd.). This equation was then used to interpolate MWi at a given intensity. Subsequently 20 row averages were taken for MWi and Inti prior to calculating: Σ(Inti)/Σ(Inti/MWi) for each lane was performed as described to determine the average telomere length.

### Chromosome-end-specific alignments

We used the recently published Telometer analysis pipeline for chromosome-end-specific alignments^41^. Initially fastq files were aligned and indexed with minimap2 against the combined T2T-CHM3, appended subtelomere reference sequence (available also on the Github) as recommended. Telomere reads were then identified with a minimum read length of 1000 bp and telomere repeat gap tolerance of 20 bp (“-m 1000, -g 20”) using V1.0 of the published pipeline available at https://github.com/santiago-es/Telometer. Telomere mapping was evaluated as the fraction of telomere reads with MAPQ filter > 30 out of total reads without mapping.

### TVR Cluster analysis and Nanopore telomere length measurement

TVR clustering and telomere length measurements were performed on a per-sample basis with Telogator2 (https://github.com/zstephens/telogator2) using default settings for oxford nanopore sequencing except for the for the following adjusted parameters to account for the digestion step in our telomere enrichment protocol: minimum read length (-l) was lowered to 1000bp, minimum length of detected subtelomeric sequence (--filt-sub) was lowered to 200bp, and the minimum number of reads for defining clusters (-n) was set per-sample by identifying the number of reads passing the previous telomeric thresholds and assuming a 5-fold difference in read depth for identified clusters (# telomeric reads/92 chromosome ends/5) or a minimum of 5 reads, whichever was higher.

Following Telogator2 clustering, inter-sample cluster correlation was calculated by pairwise alignment of TVR consensus sequences across all samples sequenced. As reads without TVR sequences could not be compared in this manner, they were excluded from the analysis. Haplotype clusters were considered a “match” if the TVR sequence similarity exceeded a Levenshtein ratio of 0.85, given the inherent sequencing error of nanopore sequencing and the repetitive nature of telomere sequences. This threshold was experimentally determined to maximize the number of aligned haplotypes between family members while minimizing cross-family TVR matching. Because TVR sequences often contain large stretches of canonical telomeric repeats, the total telomere length for individual reads was calculated by the length of the TVR + the length of the terminal canonical telomeric repeats. All results of this clustering and individual read telomere length calculation are included in Supplementary Table 2.

Results were plotted using matplotlib and the seaborn library using custom python scripts.

TVR sequences were encoded as reported for Telogator2^43^ and plotted using the following color schema:

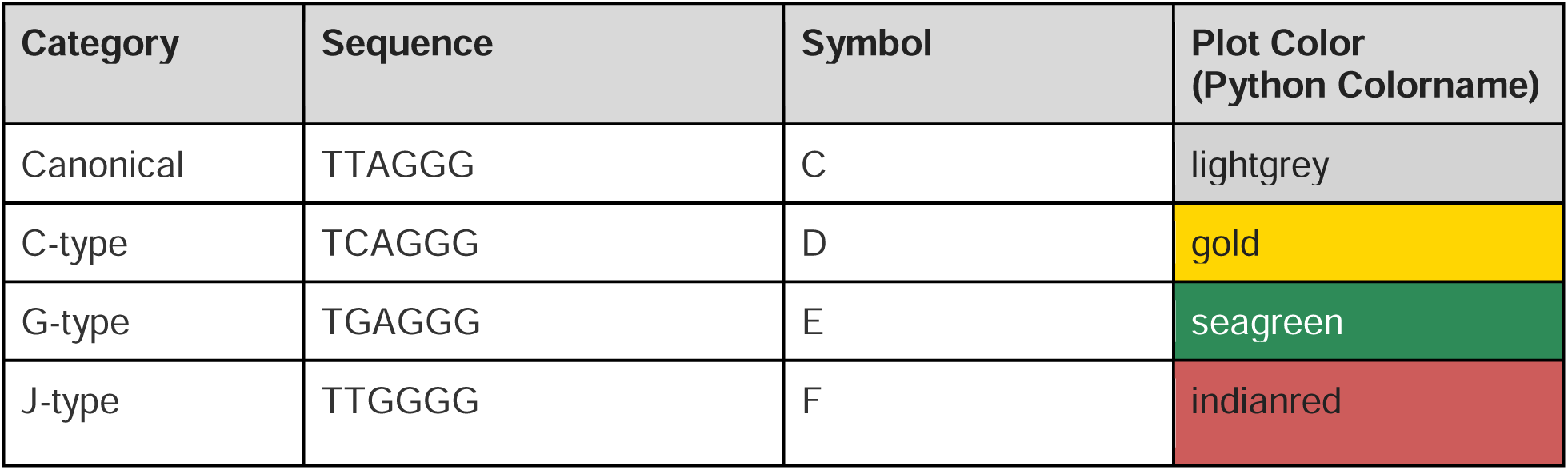
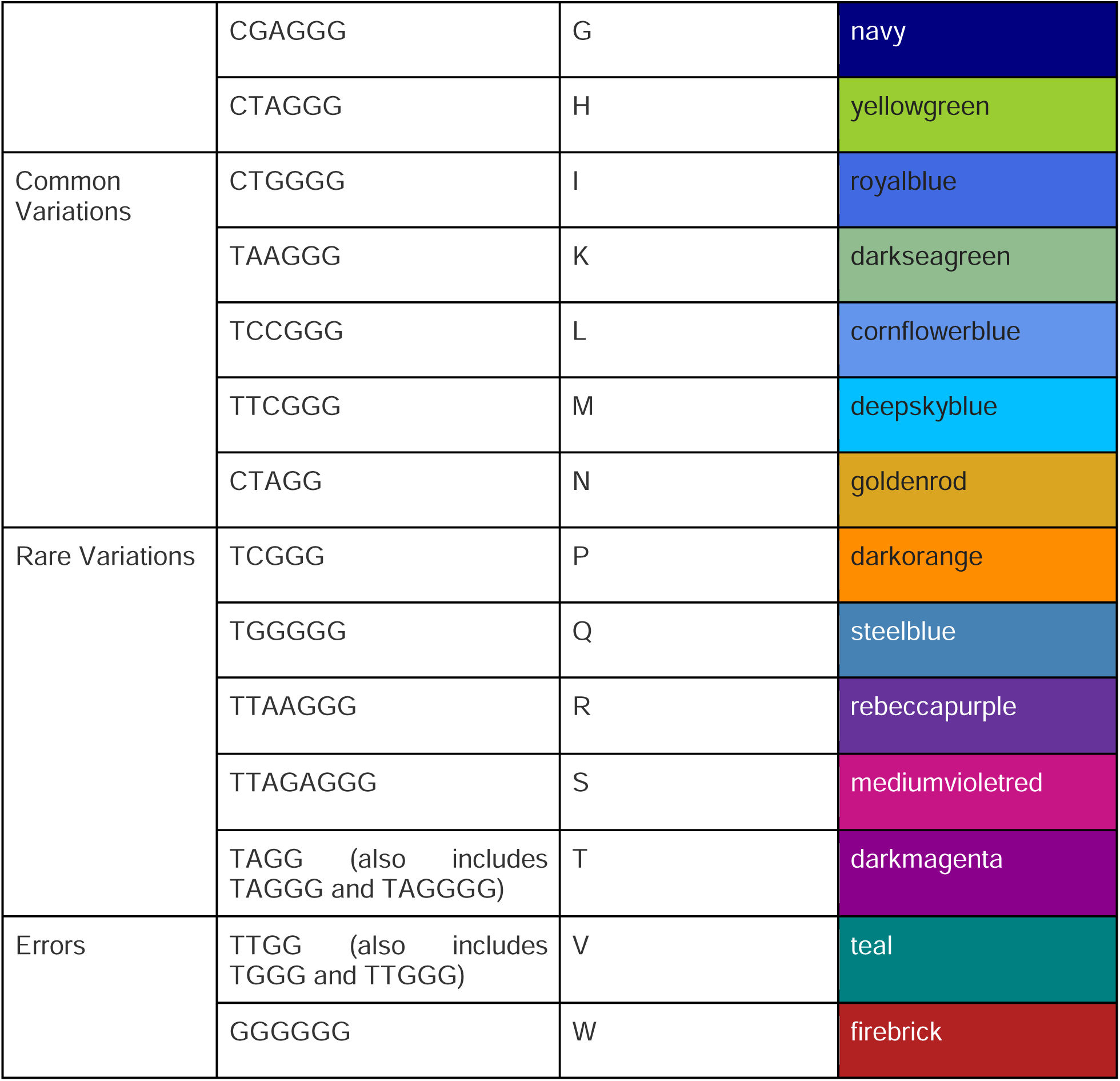

**Supplementary Table 1.**
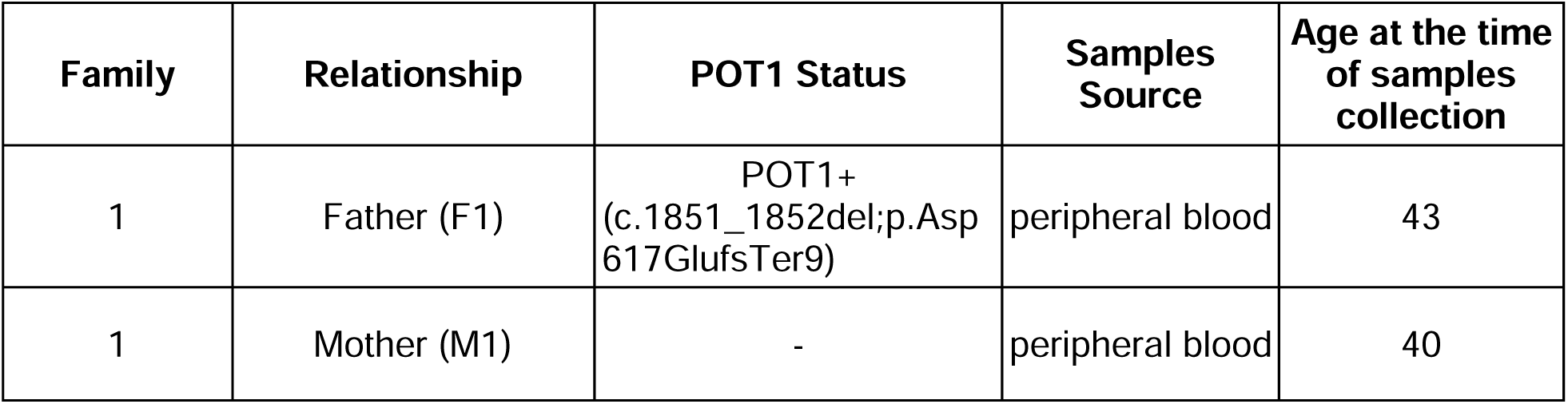
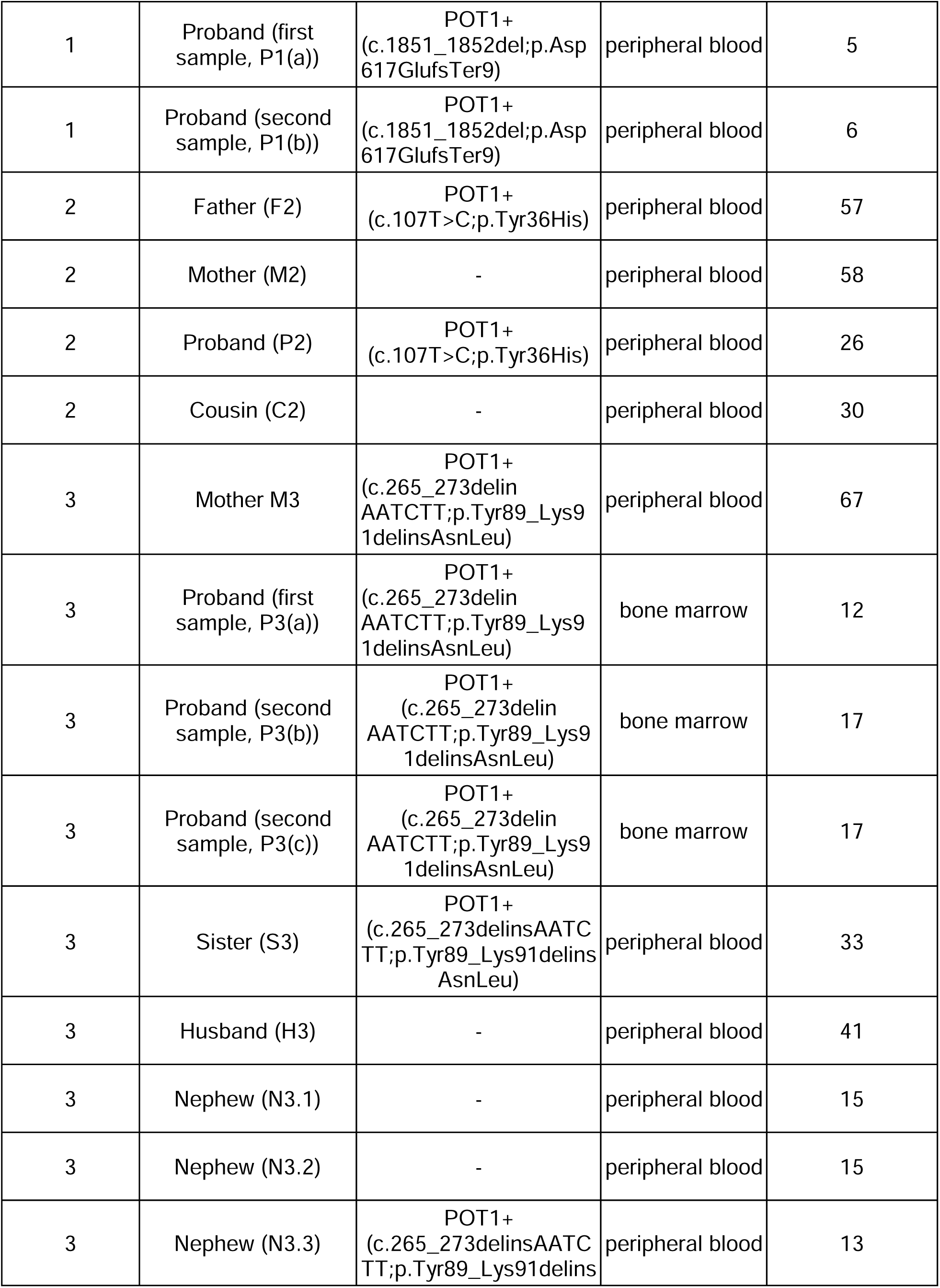
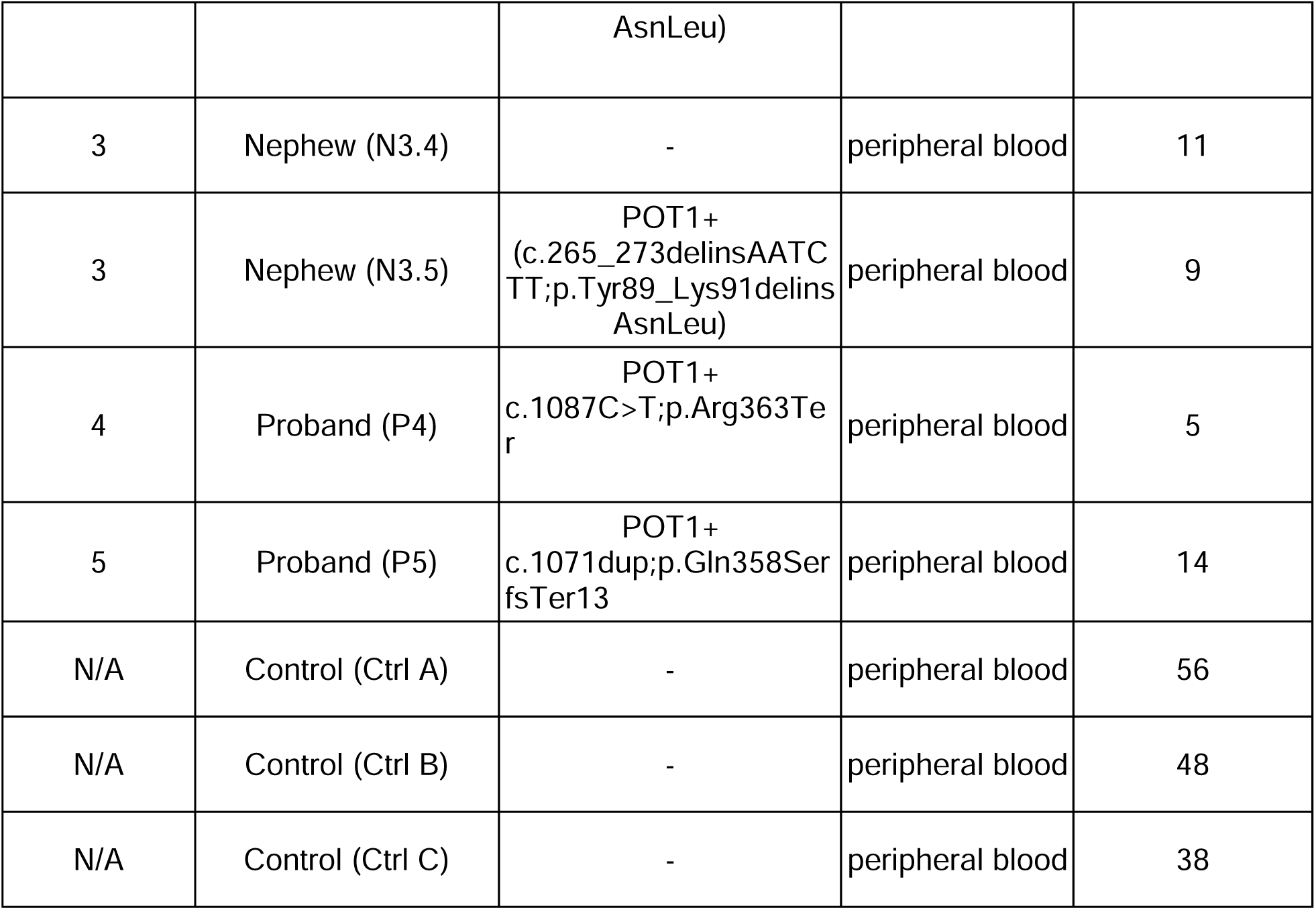

**Supplementary Table 2.** Results of telomere TVR cluster correlation, showing the consensus TVR sequence and mean levenshtein ratio for cluster correlation as well as individual sample measurements. Nan values indicate that a given TVR cluster was not identified within that sample.

**Supplementary Table 3.**
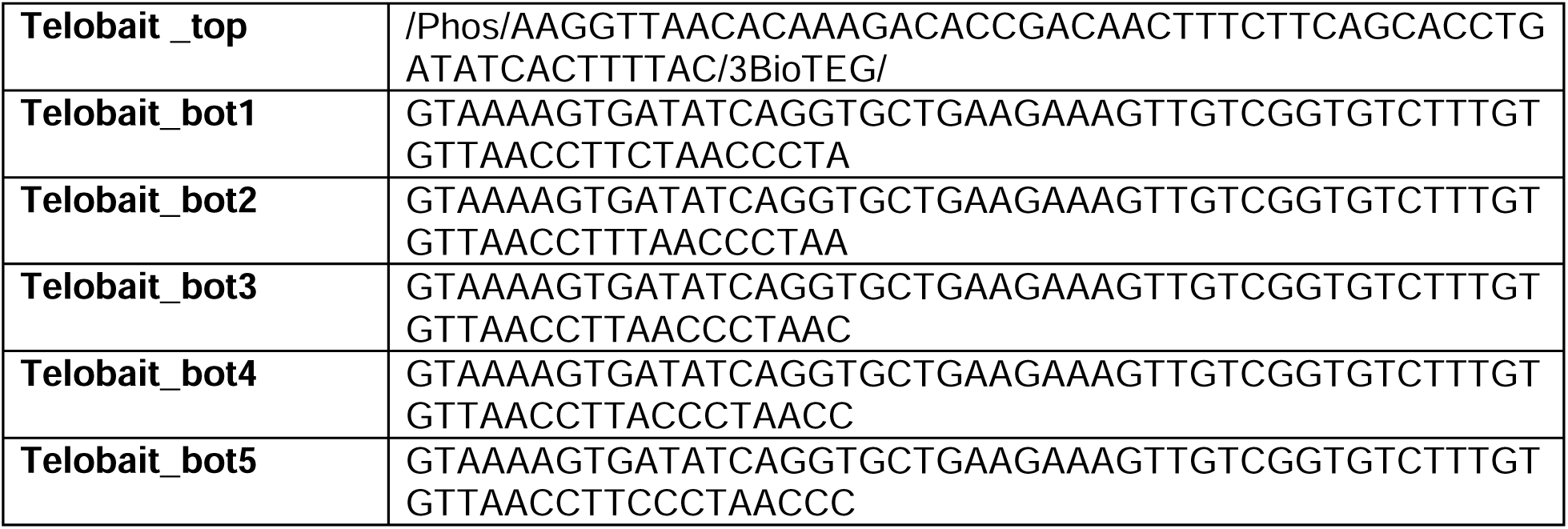
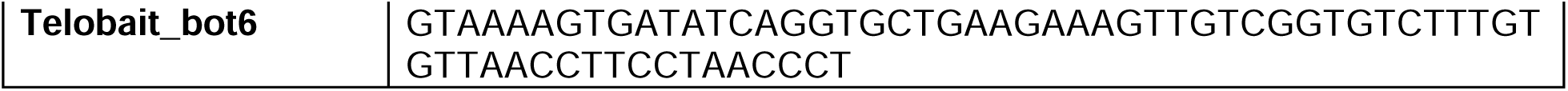
Telobait sequence for oligo synthesis ordering (5’ - 3’). Note the phosphorylation and 3’-Biotin-TEG modification for Telobait_GO_top. Telobait_GO_bot1-6 sequences are not modified. All Telobait sequences were ordered from IDT with HPLC purification.

**Supplementary Figure 1:** Telomere library preparation for nanopore sequencing Schematic of telomere enrichment and nanopore library preparation protocol

**Supplementary Figure 2:** Comparison of telomere restriction fragment length analysis to nanopore sequencing

(A) Telomere restriction fragment (TRF) length analysis of healthy unrelated controls compared against corresponding POT1 patient DNA samples used in this study from Family (1), (2), (3) as shown in Figure 1A. Mean telomere length determined by densitometry analysis of TRF are indicated at the bottom of each lane alongside ethidium bromide (EtBr) staining. (B) Comparison of telomere length measurements by TRF and nanopore sequencing. TRF mean telomere lengths from (A) versus 75th percentile bulk nanopore sequencing telomere lengths. For B, calculated values of 75pct telomere length (ATL = adjusted telomere length. ATL = TL_len+tvr_len) from Nanopore sequencing shown in Figure 1A in Supplementary Data 2 and mean telomere length of TRF calculations are shown in Supplementary Data 3.

**Supplementary Figure 3:** Comparison of sample consistency between sequencing rounds

Bulk telomere length measurements for all samples split by sequencing round. Each sequencing round refers to independent library preparation and subsequent sequencing from the same starting DNA sample. Median and interquartile range are indicated.

**Supplementary Figure 4:** Chromosome-end specific telomere alignment to standard reference genome using Telometer^38^

Chromosome-end specific telomere length measurements for Proband 3 samples a (top panel) and b (bottom panel) with p-arm alignment indicated in red and q-arm alignment shown in blue. Sample median displayed for each aligned sample. Due to the high heterogeneity of subtelomeric regions and loss of distal unique sequences during enrichment, on average, fewer than 61% of telomeric reads mapped confidently to unique chromosome ends with a MAPQ > 30 filter. Moreover, the distribution of aligned reads was skewed, with up to 430-fold more reads mapping to the most represented telomere compared to the least. These findings are consistent with results from alternative pipelines, underscoring the inherent challenges of chromosome-end assignment without a sample-specific telomere-to-telomere reference genome^38–42^. Moreover, chromosome-end specificity could in principle identify individual telomere pairs, but it does not resolve maternal vs. paternal inheritance or the distinct dynamics of allelic telomeres.

**Supplementary Figure 5:** Correlation of cluster-specific telomere length

(A) 75th percentile telomere length for each TVR cluster identified in both independently collected samples for Proband 1 and Proband 3. (B) Correlation of cluster-specific telomere length for each TVR cluster for independent sequencing runs on the same DNA sample for Proband 3. Ordinary least squares linear regression was performed using the statsmodels.api OLS module.

**Supplementary Figure 6:** Significant enrichment of carrier-inherited alleles among the longest telomeres

Intra-sample telomere rank order (A) and 75th percentile telomere length (B) plotted for each telomere allele, split by inheritance from the carrier (red) or non-carrier (blue, grey if non-carrier presumed but no non-carrier parental sample available) for the following samples: Proband 1 (left), Proband 2 (middle-left), Proband 3 (middle-right) and Sister 3 (right). Sample comparison performed using Welch’s t-test (** = p-value<0.01, *** = p-value<0.001,**** = p-value<0.0001)

**Supplementary Figure 7:** Correlation of Proband and parental absolute telomere lengthlength Proband (x-axis) and parental (y-axis) 75th percentile telomere length for shared alleles inherited from the carrier (red) and non-carrier (blue) parents for Proband 1 (A) and Proband 2

(B). Ordinary least squares linear regression was performed independently for carrier and non-carrier alleles using the statsmodels.api OLS module.

**Supplementary Figure 8:** Proband 2 shows significantly longer shared telomere length than their non-carrier Cousin 2

Comparison of 75th percentile telomere length between Proband 2 and Cousin 2, subset to include only shared TVR haplotypes. Sample comparison performed using Welch’s t-test (**** = p-value<0.0001)

## References

1. Shi, J. et al. Rare missense variants in POT1 predispose to familial cutaneous malignant melanoma. Nat. Genet. 46, 482–486 (2014).

2. Robles-Espinoza, C. D. et al. POT1 loss-of-function variants predispose to familial melanoma. Nat. Genet. 46, 478–481 (2014).

3. Schratz, K. E. et al. T cell immune deficiency rather than chromosome instability predisposes patients with short telomere syndromes to squamous cancers. Cancer Cell 41, 807–817.e6 (2023).

4. DeBoy, E. A. et al. Familial Clonal Hematopoiesis in a Long Telomere Syndrome. N. Engl. J. Med. 388, 2422–2433 (2023).

5. DeBoy, E. A. et al. Telomere-lengthening germline variants predispose to a syndromic papillary thyroid cancer subtype. Am. J. Hum. Genet. 111, 1114–1124 (2024).

6. Calvete, O. et al. The wide spectrum of POT1 gene variants correlates with multiple cancer types. Eur. J. Hum. Genet. 25, 1278–1281 (2017).

7. Jalali, A. et al. POT1 Regulates Proliferation and Confers Sexual Dimorphism in Glioma. Cancer Res. 81, 2703–2713 (2021).

8. Shen, E. et al. POT1 mutation spectrum in tumour types commonly diagnosed among POT1-associated hereditary cancer syndrome families. J. Med. Genet. 57, 664–670 (2020).

9. Ramsay, A. J. et al. POT1 mutations cause telomere dysfunction in chronic lymphocytic leukemia. Nat. Genet. 45, 526–530 (2013).

10. McMaster, M. L. et al. Germline Mutations in Protection of Telomeres 1 in Two Families with Hodgkin Lymphoma. Br. J. Haematol. 181, 372–377 (2018).

11. Gilene, S., Knapke, S., Leino, D., Roy, S. & Raskin, S. A novel POT1-TPD presentation: A germline pathogenic POT1 variant discovered in a patient with newly diagnosed posterior fossa ependymoma. Cancer Genet. **292–293**, 38–43 (2025).

12. Wu, Y., Poulos, R. C. & Reddel, R. R. Role of POT1 in Human Cancer. Cancers 12, 2739 (2020).

13. Kim, W.-T. et al. Cancer-associated POT1 mutations lead to telomere elongation without induction of a DNA damage response. EMBO J. 40, e107346 (2021).

14. Baumann, P. & Cech, T. R. Pot1, the Putative Telomere End-Binding Protein in Fission Yeast and Humans. Science 292, 1171–1175 (2001).

15. Loayza, D. & de Lange, T. POT1 as a terminal transducer of TRF1 telomere length control. Nature 423, 1013–1018 (2003).

16. Lei, M., Podell, E. R. & Cech, T. R. Structure of human POT1 bound to telomeric single-stranded DNA provides a model for chromosome end-protection. Nat. Struct. Mol. Biol. 11, 1223–1229 (2004).

17. Denchi, E. L. & de Lange, T. Protection of telomeres through independent control of ATM and ATR by TRF2 and POT1. Nature 448, 1068–1071 (2007).

18. Hockemeyer, D., Sfeir, A. J., Shay, J. W., Wright, W. E. & de Lange, T. POT1 protects telomeres from a transient DNA damage response and determines how human chromosomes end. EMBO J. 24, 2667–2678 (2005).

19. Zaug, A. J., Podell, E. R. & Cech, T. R. Human POT1 disrupts telomeric G-quadruplexes allowing telomerase extension in vitro. Proc. Natl. Acad. Sci. U. S. A. 102, 10864–10869 (2005).

20. Martin, A. et al. Active telomere elongation by a subclass of cancer-associated POT1 mutations. Genes Dev. 39, 445–462 (2025).

21. Takai, H. et al. A POT1 mutation implicates defective telomere end fill-in and telomere truncations in Coats plus. Genes Dev. 30, 812–826 (2016).

22. Tummala, H. et al. The evolving genetic landscape of telomere biology disorder dyskeratosis congenita. EMBO Mol. Med. 16, 2560–2582 (2024).

23. Kelich, J. et al. Telomere dysfunction implicates POT1 in patients with idiopathic pulmonary fibrosis. J. Exp. Med. 219, e20211681 (2022).

24. Savage, S. A., Bertuch, A. A. & Team Telomere and the Clinical Care Consortium for Telomere-Associated Ailments (CCCTAA). Different phenotypes with different endings—Telomere biology disorders and cancer predisposition with long telomeres. Br. J. Haematol.

25. Revy, P., Kannengiesser, C. & Bertuch, A. A. Genetics of human telomere biology disorders. Nat. Rev. Genet. 24, 86–108 (2023).

26. Armanios, M. The Role of Telomeres in Human Disease. Annu. Rev. Genomics Hum. Genet. 23, 363–381 (2022).

27. Vulliamy, T. et al. Disease anticipation is associated with progressive telomere shortening in families with dyskeratosis congenita due to mutations in TERC. Nat. Genet. 36, 447–449 (2004).

28. Armanios, M. et al. Haploinsufficiency of telomerase reverse transcriptase leads to anticipation in autosomal dominant dyskeratosis congenita. Proc. Natl. Acad. Sci. 102, 15960–15964 (2005).

29. Hemann, M. T., Strong, M. A., Hao, L. Y. & Greider, C. W. The shortest telomere, not average telomere length, is critical for cell viability and chromosome stability. Cell 107, 67–77 (2001).

30. Zou, Y., Sfeir, A., Gryaznov, S. M., Shay, J. W. & Wright, W. E. Does a sentinel or a subset of short telomeres determine replicative senescence? Mol. Biol. Cell 15, 3709–3718 (2004).

31. Kaul, Z., Cesare, A. J., Huschtscha, L. I., Neumann, A. A. & Reddel, R. R. Five dysfunctional telomeres predict onset of senescence in human cells. EMBO Rep. 13, 52–59 (2011).

32. Turner, S. & Hartshorne, G. M. Telomere lengths in human pronuclei, oocytes and spermatozoa. Mol. Hum. Reprod. 19, 510–518 (2013).

33. Nordfjäll, K., Larefalk, A., Lindgren, P., Holmberg, D. & Roos, G. Telomere length and heredity: Indications of paternal inheritance. Proc. Natl. Acad. Sci. U. S. A. 102, 16374–16378 (2005).

34. Njajou, O. T. et al. Telomere length is paternally inherited and is associated with parental lifespan. Proc. Natl. Acad. Sci. U. S. A. 104, 12135–12139 (2007).

35. Asghar, M., Bensch, S., Tarka, M., Hansson, B. & Hasselquist, D. Maternal and genetic factors determine early life telomere length. Proc. R. Soc. B Biol. Sci. 282, 20142263 (2015).

36. Kimura, M. et al. Offspring’s leukocyte telomere length, paternal age, and telomere elongation in sperm. PLoS Genet. 4, e37 (2008).

37. Baird, D. M., Rowson, J., Wynford-Thomas, D. & Kipling, D. Extensive allelic variation and ultrashort telomeres in senescent human cells. Nat. Genet. 33, 203–207 (2003).

38. Lim, T. L. et al. Germline POT1 variants can predispose to myeloid and lymphoid neoplasms. Leukemia 36, 283–287 (2022).

39. Nathan, V. et al. Loss-of-function variants in POT1 predispose to uveal melanoma. J. Med. Genet. 58, 234–236 (2021).

40. Chen, C. et al. Structural insights into POT1-TPP1 interaction and POT1 C-terminal mutations in human cancer. Nat. Commun. 8, 14929 (2017).

41. Sanchez, S. E. et al. Digital telomere measurement by long-read sequencing distinguishes healthy aging from disease. Nat. Commun. 15, 5148 (2024).

42. Smoom, R. et al. Telomouse—a mouse model with human-length telomeres generated by a single amino acid change in RTEL1. Nat. Commun. 14, 6708 (2023).

43. Karimian, K. et al. Human telomere length is chromosome end-specific and conserved across individuals. Science 384, 533–539 (2024).

44. Schmidt, T. T. et al. High resolution long-read telomere sequencing reveals dynamic mechanisms in aging and cancer. https://doi.org/10.1101/2023.11.28.569082 (2023) doi:10.1101/2023.11.28.569082.

45. Smoom, R., Lichtental, D., Kaestner, K. H. & Tzfati, Y. The house mouse maintains constant telomere length throughout life. Nucleic Acids Res. 53, gkaf830 (2025).

46. Stephens, Z. & Kocher, J.-P. Characterization of telomere variant repeats using long reads enables allele-specific telomere length estimation. BMC Bioinformatics 25, 194 (2024).

47. Colgin, L. M., Baran, K., Baumann, P., Cech, T. R. & Reddel, R. R. Human POT1 facilitates telomere elongation by telomerase. Curr. Biol. CB 13, 942–946 (2003).

48. Lin, T. T. et al. Telomere dysfunction and fusion during the progression of chronic lymphocytic leukemia: evidence for a telomere crisis. Blood 116, 1899–1907 (2010).

49. Guièze, R. et al. Telomere status in chronic lymphocytic leukemia with TP53 disruption. Oncotarget 7, 56976–56985 (2016).

50. Telomeres Mendelian Randomization Collaboration et al. Association Between Telomere Length and Risk of Cancer and Non-Neoplastic Diseases: A Mendelian Randomization Study. JAMA Oncol. 3, 636–651 (2017).

51. Rode, L., Nordestgaard, B. G. & Bojesen, S. E. Long telomeres and cancer risk among 950568 individuals from the general population. Int. J. Epidemiol. 45, 1634–1643 (2016).

52. McNally, E. J., Luncsford, P. J. & Armanios, M. Long telomeres and cancer risk: the price of cellular immortality. J. Clin. Invest. 129, 3474–3481 (2019).

53. Nassour, J., Schmidt, T. T. & Karlseder, J. Telomeres and Cancer: Resolving the Paradox. Annu. Rev. Cancer Biol. 5, 59–77 (2021).

54. Tham, C.-Y. et al. High-throughput telomere length measurement at nucleotide resolution using the PacBio high fidelity sequencing platform. Nat. Commun. 14, 281 (2023).

55. Mender, I. & Shay, J. W. Telomere Restriction Fragment (TRF) Analysis. Bio-Protoc. 5, e1658 (2015).

56. Rueden, C. T. et al. ImageJ2: ImageJ for the next generation of scientific image data. BMC Bioinformatics 18, 529 (2017).

