## Supplemental Figures for "Telomere length of both parents contributes to heritable POT1 cancer-predisposition syndrome"

### Martin et al., Supplementary Figure 1

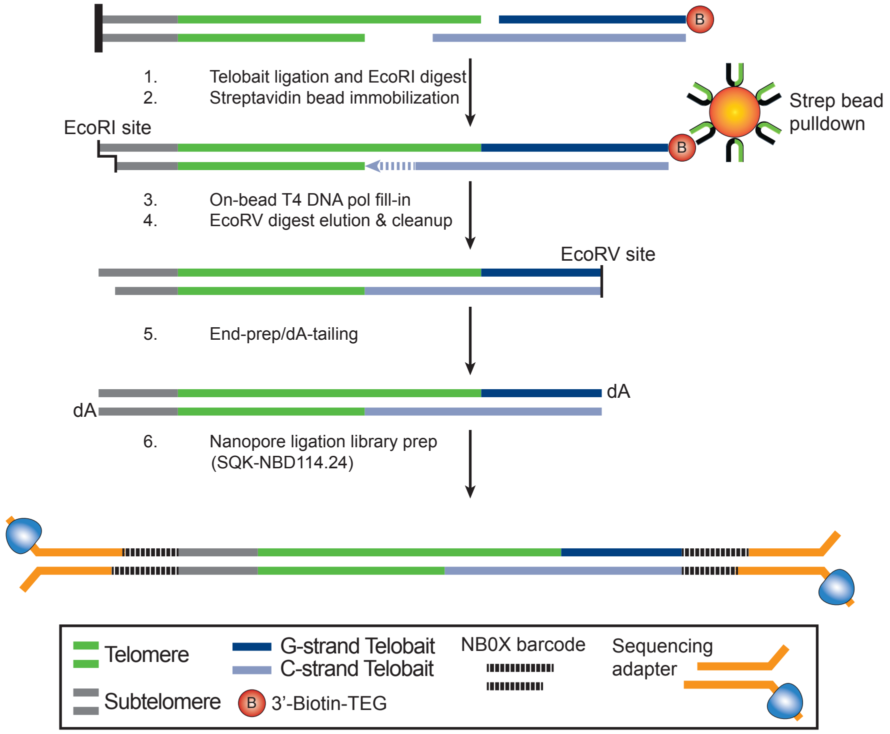

### Martin et al., Supplementary Figure 2

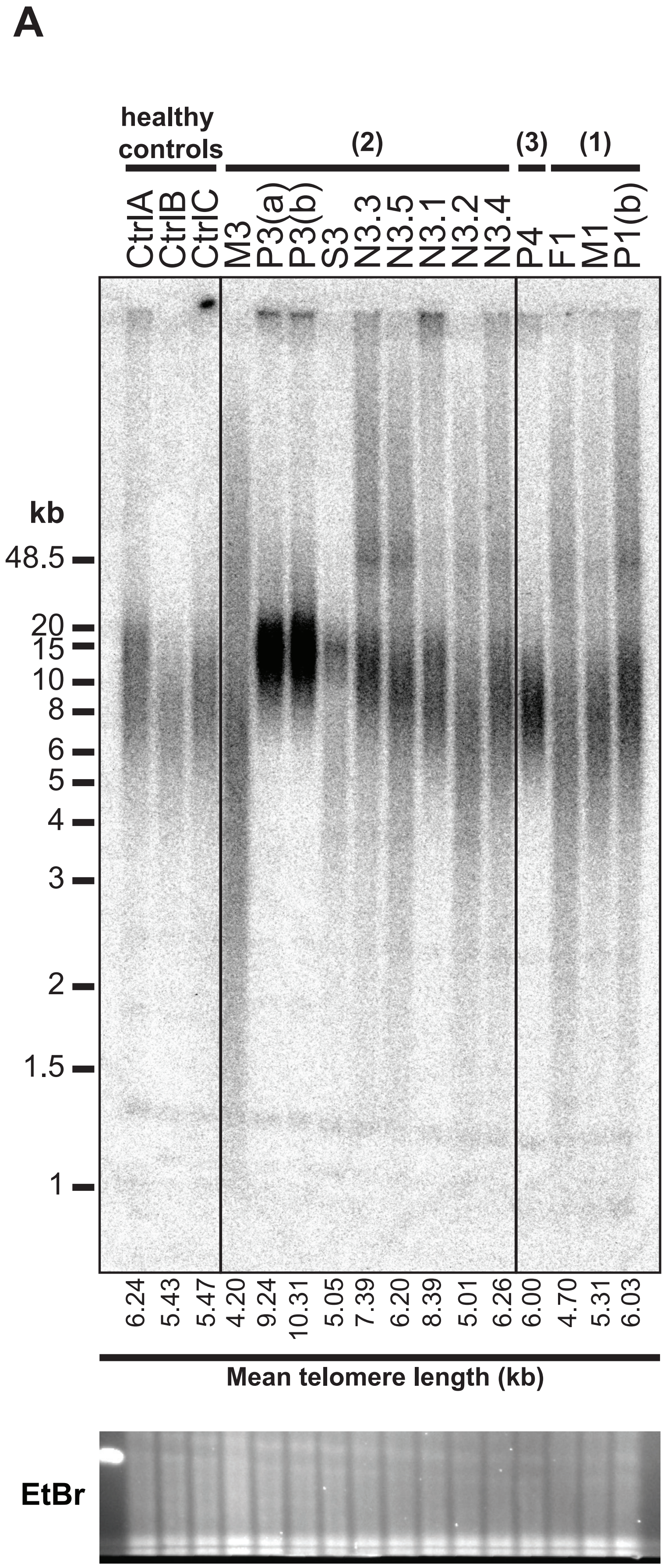


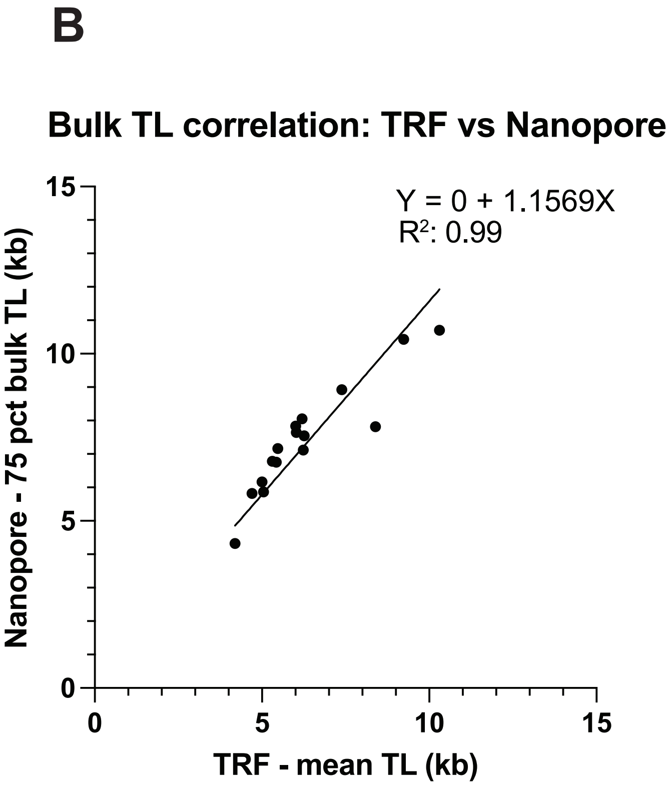

### Martin et al., Supplementary Figure 3

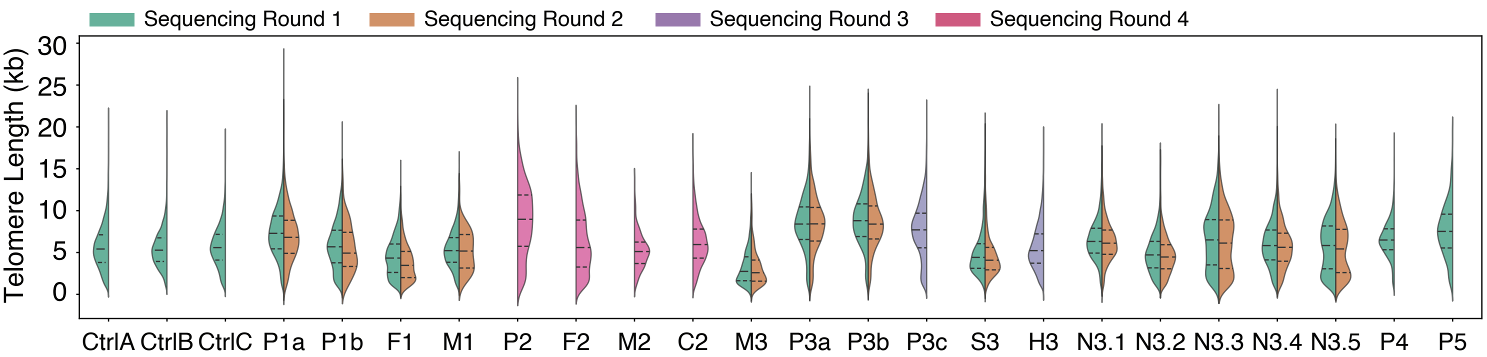

### Martin et al., Supplementary Figure 4

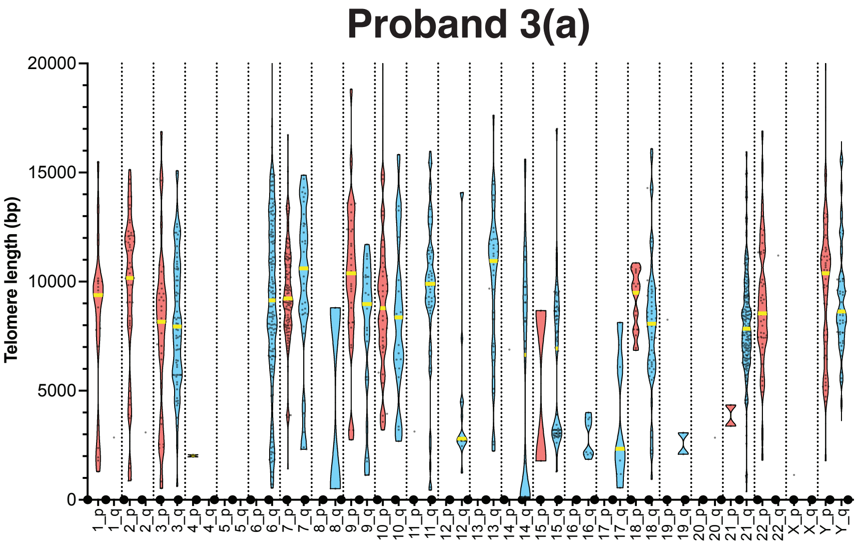


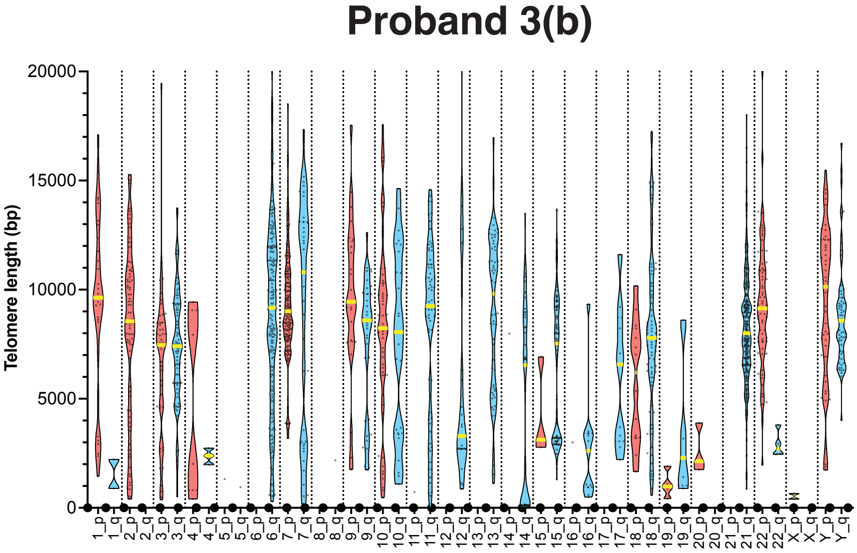

### Martin et al., Supplementary Figure 5

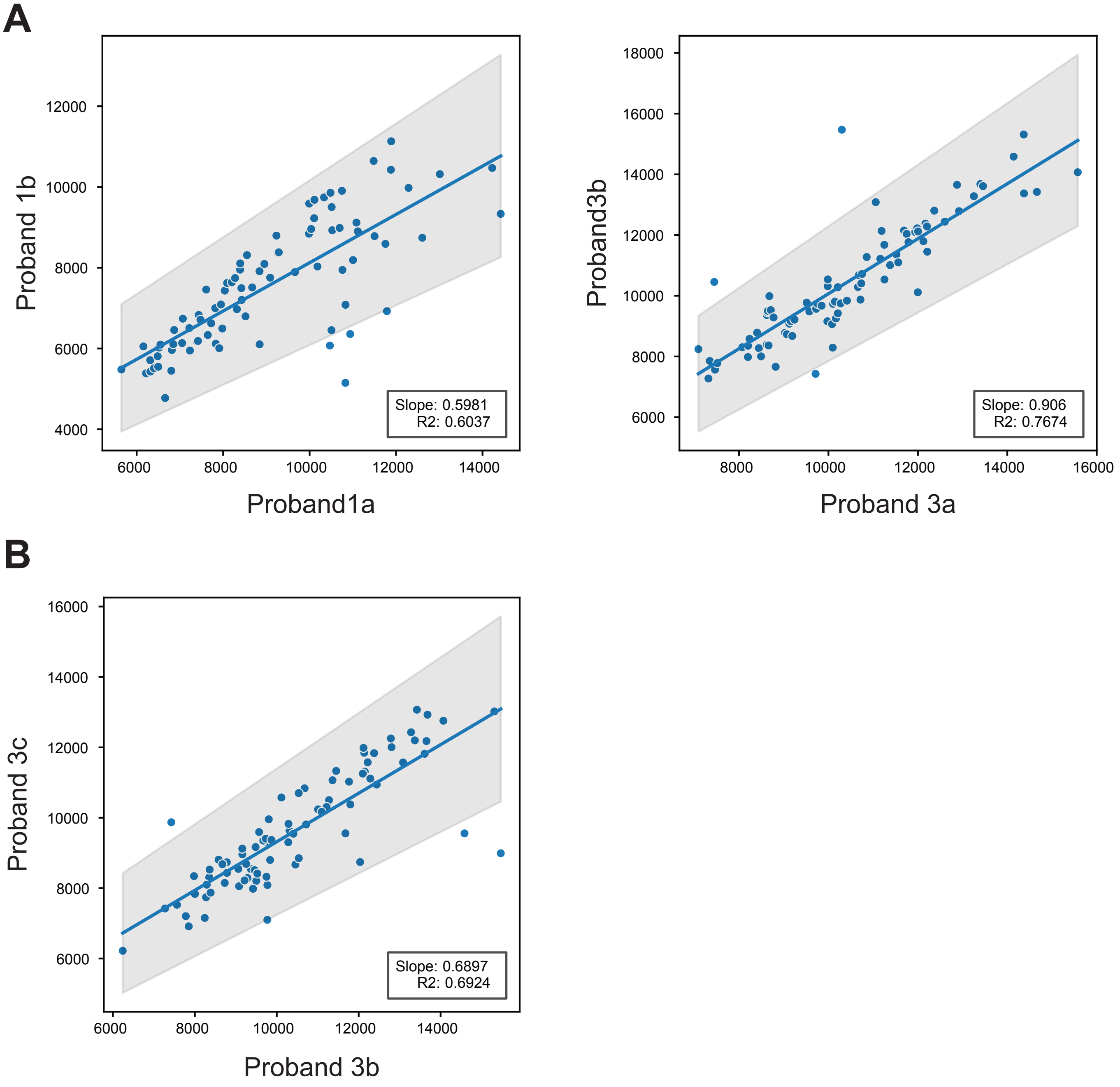

### Martin et al., Supplementary Figure 6

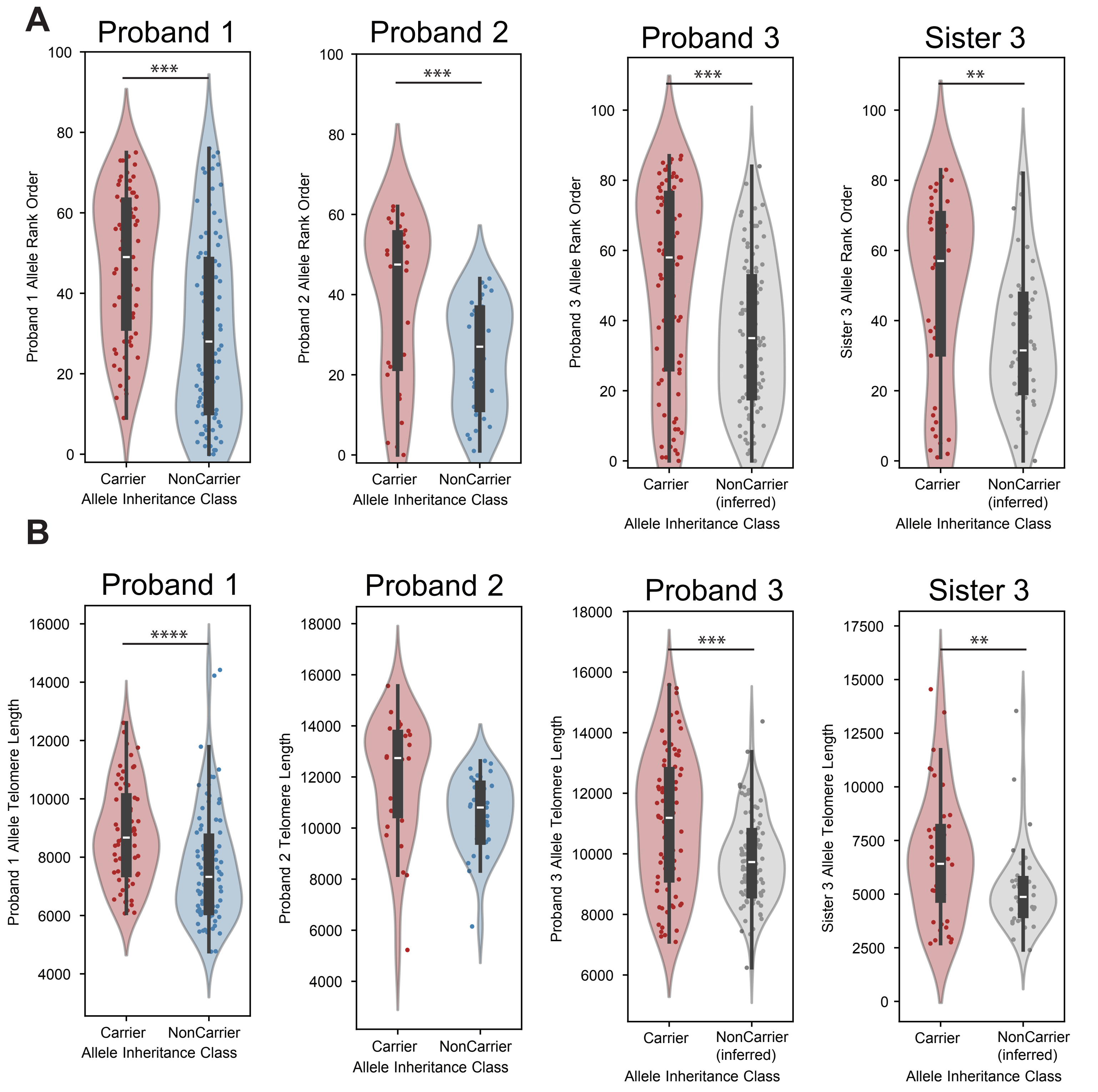

### Martin et al, Supplementary Figure 7

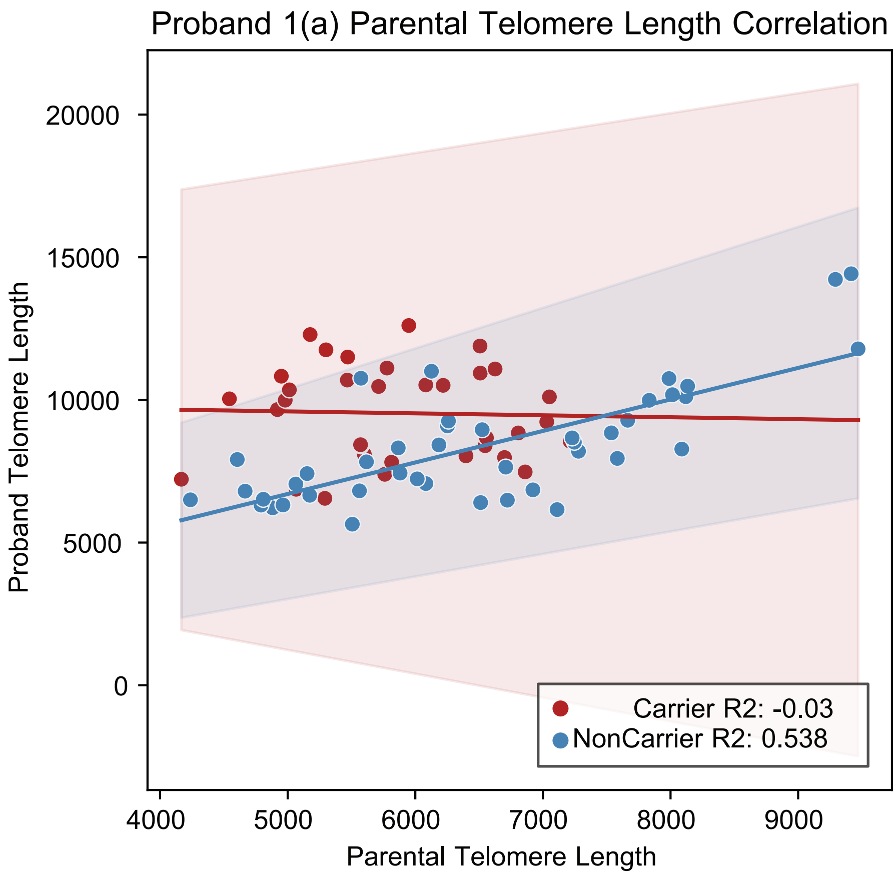


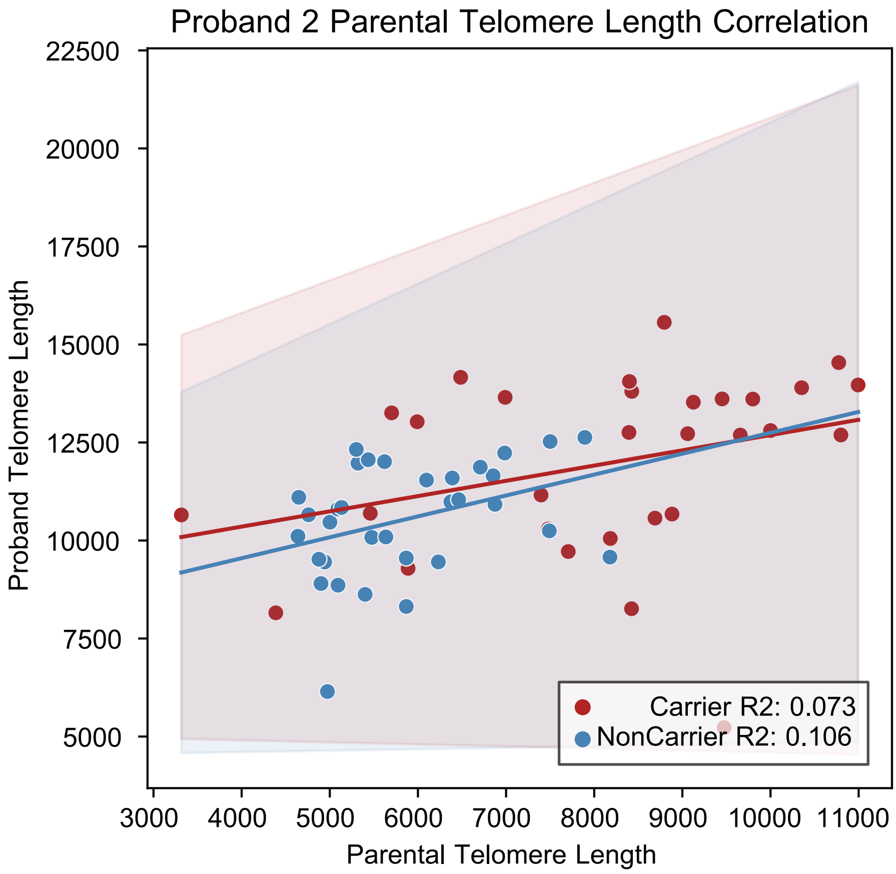

### Martin et al, Supplementary Figure 8

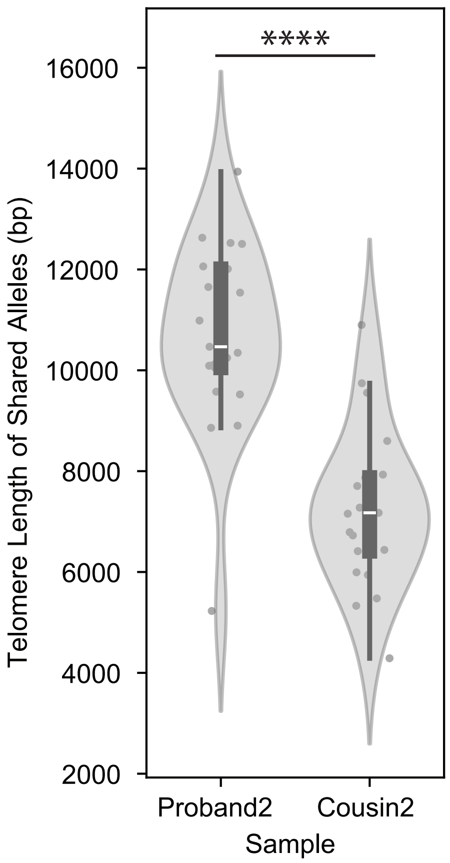
